# *Atrx* deletion in neurons leads to sexually-dimorphic dysregulation of miR-137 and spatial learning and memory deficits

**DOI:** 10.1101/606442

**Authors:** Renee J. Tamming, Vanessa Dumeaux, Luana Langlois, Jacob Ellegood, Lily R. Qiu, Yan Jiang, Jason P. Lerch, Nathalie G. Bérubé

## Abstract

Mutations in the ATRX chromatin remodeler are associated with syndromic and non-syndromic intellectual disability. Emerging evidence points to key roles for ATRX in preserving neuroprogenitor cell genomic stability, whereas ATRX function in differentiated neurons and memory processes are still unresolved. Here, we show that *Atrx* deletion in mouse forebrain glutamatergic neurons causes distinct hippocampal structural defects identified by magnetic resonance imaging. Ultrastructural analysis revealed fewer presynaptic vesicles and an enlarged postsynaptic area at CA1 apical dendrite-axon junctions. These synaptic defects are associated with impaired long-term contextual memory in male, but not female mice. Mechanistically, we identify ATRX-dependent and sex-specific alterations in synaptic gene expression linked to Mir137 levels, a known regulator of presynaptic processes and spatial memory. We conclude that ablation of *Atrx* in excitatory forebrain neurons leads to sexually dimorphic outcomes on miR-137 and on spatial memory, identifying a promising therapeutic target for neurological disorders caused by ATRX dysfunction.

**Summary statement:** Ablation of the ATRX chromatin remodeler specifically in forebrain excitatory neurons of mice causes male-specific deficits in long-term spatial memory associated with miR-137 overexpression, transcriptional changes and structural alterations corresponding to pre- and post-synaptic abnormalities.

## Introduction

Alpha-thalassemia X-linked intellectual disability syndrome, or ATR-X syndrome, is a rare congenital X-linked disorder resulting in moderate to severe intellectual disability (ID), developmental delay, microcephaly, hypomyelination, and a mild form of alpha-thalassemia [OMIM: 301040]^1^. In a recent study of approximately 1000 individuals with ID, *ATRX* mutations were identified as one of the most frequent cause of non-syndromic ID^2^, emphasizing a key requirement for this gene in cognitive processes. ATRX-related ID arises from hypomorphic mutations in the *ATRX* gene, most commonly in the highly conserved ATRX/DNMT3/DNMT3L (ADD) and Switch/Sucrose non-fermenting (SWI/SNF) domains^3,4^. The former targets ATRX to chromatin by means of a histone reader domain that recognizes specific histone tail modifications^5^, and the latter confers ATPase activity and is critical for its chromatin remodeling activity^6,7^.

ATRX, in a complex with the histone chaperone DAXX, promotes the deposition of the histone variant H3.3 at heterochromatic domains including telomeres and pericentromeres^8,9^. However, ATRX is also required for H3.3 deposition within the gene body of a subset of G-rich genes, presumably to reduce G-quadruplex formation and promote transcriptional elongation^10^. ATRX is also required for the postnatal suppression of a network of imprinted genes in the neonatal brain by promoting long range chromatin interactions via CTCF and cohesin^11^.

In mice, germline deletion of *Atrx* results in embryonic lethality^12^ while conditional deletion of *Atrx* in neuroprogenitors leads to excessive DNA damage caused by DNA replication stress and subsequent Tp53-dependent apoptosis^13,14^. Mice with deletion of exon 2 of *Atrx* (*Atrx*^*ΔE2*^) were generated that result in global reduction of *Atrx* expression. These mice are viable and exhibit impaired novel object recognition memory, spatial memory in the Barnes maze, and contextual fear memory^15^. Some of the molecular defects identified in these mice included decreased activation of CaMKII and the AMPA receptor in the hippocampus as well as decreased spine density in the medial prefrontal cortex, and altered DNA methylation and increased expression of *Xlr3b* in neurons^16^. Our group also reported similar behavioural impairments in female mice exhibiting mosaic expression of ATRX in the central nervous system^17^. However, the contribution of different cell types and sex difference to behavioural abnormalities have not yet been resolved.

To start addressing these questions, we deleted *Atrx* specifically in glutamatergic forebrain neurons in male and female mice. This approach bypasses deleterious effects of ATRX loss of function that we previously observed during brain development caused by replication stress in proliferating neuroprogenitors^12,13^. A comprehensive analysis of these mice reveals that ATRX promotes long-term spatial learning and memory associated with morphological and synaptic ultrastructural changes in the hippocampus. We show that female mice lacking ATRX in neurons are protected from spatial learning and memory defects and identify sex-specific effects of ATRX loss on the expression of synaptic genes and miR-137. Overall, we identify a novel sex-specific function for ATRX in neurons in the regulation of long-term spatial memory associated with abnormal synapse ultrastructure.

## Methods

### Animal care and husbandry

Mice were exposed to a 12-hour-light/12-hour-dark cycle and with water and chow *ad libitum*. The *Atrx*^*loxP*^ mice have been described previously^18^. *Atrx*^*loxP*^ mice were mated with C57BL/6 mice expressing Cre recombinase under the control of the *αCaMKII* gene promoter^19^. The progeny includes hemizygous male mice that produce no ATRX protein in forebrain excitatory neurons (*Atrx*-cKO). The *Atrx*-cKO males were mated to *Atrx*^*loxP*^ females to yield homozygous deletion of *Atrx* in female mice (*Atrx*-cKO^Fem^). Male and female littermate floxed mice lacking the Cre allele were used as controls (Ctrl; Ctrl^Fem^). Genotyping of tail biopsies for the presence of the floxed and Cre alleles was performed as described previously^18^. All procedures involving animals were conducted in accordance with the regulations of the Animals for Research Act of the province of Ontario and approved by the University of Western Ontario Animal Care and Use Committee (2017-048). Behavioural assessments started with less demanding tasks (open field tests, elevated plus maze) to more demanding ones (Morris water maze). ARRIVE guidelines were followed: mouse groups were randomized, experimenters were blind to the genotypes, and software-based analysis was used to score mouse performance in all the tasks. All behavioural experiments were performed between 9:00 AM and 4:00 PM.

### Immunofluorescence staining

Mice were perfused with 25mL phosphate buffered saline (PBS) followed by 25mL 4% paraformaldehyde (PFA) in PBS and the brain fixed for 24 hours in 4% PFA in PBS and cryopreserved in 30% sucrose/PBS. Brains were flash frozen in Cryomatrix (Thermo Scientific) and sectioned at 8µm thickness as described previously^13^. For immunostaining, antigen retrieval was performed by incubating slides in 10 mM sodium citrate at 95°C for 10 min. Cooled sections were washed and blocked with 10% normal goat serum (Sigma). The slides were incubated overnight in primary antibody (ATRX: 1:200, H-300 Santa Cruz Biotechnology Inc; GFAP: 1:200, Agilent Technologies, Inc; IBA1: 1:500, Wako Pure Chemical Corporation) diluted in 0.3% Triton-X/PBS. Sections were washed in 0.3% Triton-X/PBS 3× 5 min and incubated with secondary antibody (goat anti-rabbit-Alexa Fluor 594: 1:800, Life Technologies) for one hour. Sections were washed again three times for 5 min, counterstained with DAPI and mounted with SlowFade Gold (Invitrogen). All images were captured using an inverted microscope (Leica DMI 6000b) with a digital camera (Hamamatsu ORCA-ER). Openlab image software was used for manual image capture, and images were processed using the Volocity software (Demo Version 6.0.1; PerkinElmer) and Adobe Photoshop CS6 (Version 13.0). Cell counts of DAPI, GFAP, and IBA1 were performed in Adobe Photoshop. DAPI was counted per mm^2^ and GFAP and IBA1 were counted as percentages of DAPI^+^cells. One section from five pairs of Ctrl/*Atrx*-cKO was counted. Statistical significance was determined using an unpaired Student’s T-test.

### Reverse transcriptase real-time PCR (qRT-PCR)

Total RNA was isolated from control and *Atrx*-cKO frontal cortex and hippocampus using the miRVANA total RNA isolation kit (ThermoFisher) and reverse transcribed to cDNA using 1 μg RNA and SuperScript II Reverse Transcriptase (Invitrogen). Real-time PCR was performed in duplicate using gene-specific primers under the following conditions: 95°C for 10 s, 58°C for 20 s, 72°C for 30 s for 35 cycles. All data were normalized against β-actin expression levels. Primers used were as follows: *β-actin (*forward CTGTCGAGTCGCGTCCACCC, reverse ACATGCCGGAGCCGTTGTCG); *Atrx* (forward AGAAATTGAGGATGCTTCACC, reverse TGAACCTGGGGACTTCTTTG).

Total RNA was also used for reverse transcription of miRNA using the TaqMan Advanced MicroRNA reverse transcription kit (ThermoFisher). qRT-PCR was performed using the TaqMan Universal PCR Master Mix, no AmpErase (ThermoFisher) using advanced probes for miR-137-3p (mmu480924_mir), miR-34a-5p (mmu481304_mir), miR-27b-3p (mmu478270_mir), miR-485-5p (mmu482774_mir), and normalized to miR-191-5p (mmu481584_mir) under the following protocol: 95 °C for 10 min, 45 cycles of 95 °C for 15 s and 60 °C for 1 min.

### Western blot analysis

Whole cell lysates were collected in standard RIPA buffer and quantified using a Bradford assay (BioRad). Protein lysates (50µg) were loaded on a 6% SDS-PAGE gel and transferred to nitrocellulose membrane (BioRad) using a wet electroblotting system (BioRad). The membrane was blocked in 5% milk in Tris-buffered saline with 0.1% Tween-20 (SigmaAldrich) for 1 hour and incubated overnight at 4°C with primary antibody (ATRX: 1:500, H-300 Santa Cruz Biotechnology Inc.; INCENP: 1:3000, Sigma). The following day the membrane was washed and incubated in the appropriate horseradish peroxidase-conjugated secondary antibody (1:4000, Jackson ImmunoResearch Laboratories, Inc.). The membrane was rinsed briefly in enhanced chemiluminescence substrate (BioRad) before exposure on a ChemiDoc Gel Imaging System (BioRad).

### Magnetic resonance imaging

Mice were perfused with 30mL of PBS supplemented with 1µL/mL heparin (Sandoz) and 2mM ProHance (Bracco Imaging Canada) followed by 30mL 4% paraformaldehyde supplemented with 2mM ProHance. After perfusion, mice were decapitated, and the skin, cartilaginous nose tip, and lower jaw were removed. The remaining brain and skull structures were incubated in PBS supplemented with 2mM ProHance and 0.02% sodium azide for at least one month before MRI scanning^20^. A multi-channel 7.0 Tesla MRI scanner (Agilent Inc., Palo Alto, CA) was used to image the brains within their skulls. Sixteen custom-built solenoid coils were used to image the brains in parallel^21^. In order to detect volumetric changes, we used the following parameters for the MRI scan: T2-weighted, 3-D fast spin-echo sequence, with a cylindrical acquisition of k-space, a TR of 350 ms, and TEs of 12 ms per echo for 6 echoes, field-of-view equaled to 20 × 20 × 25 mm3 and matrix size equaled to 504 × 504 × 630. Our parameters output an image with 0.040 mm isotropic voxels. The total imaging time was 14 hours^22^.

### MRI Registration and Analysis

To visualize and compare any changes in the mouse brains, the images are linearly (6 followed by 12 parameter) and non-linearly registered together. Registrations were performed with a combination of mni_autoreg tools^23^ and ANTS (advanced normalization tools)^24^. All scans are then resampled with the appropriate transform and averaged to create a population atlas representing the average anatomy of the study sample. The result of the registration is to have all images deformed into alignment with each other in an unbiased fashion. For the volume measurements, this allows for the analysis of the deformations needed to take each individual mouse’s anatomy into this final atlas space, the goal being to model how the deformation fields relate to genotype^25,26^. The jacobian determinants of the deformation fields are then calculated as measures of volume at each voxel. Significant volume differences can then be calculated by warping a pre-existing classified MRI atlas onto the population atlas, which allows for the volume of 182 different segmented structures encompassing cortical lobes, large white matter structures (i.e. corpus callosum), ventricles, cerebellum, brain stem, and olfactory bulbs ^27-31^ to be assessed in all brains. Further, these measurements can be examined on a voxel-wise basis to localize the differences found within regions or across the brain. Multiple comparisons in this study were controlled for using the False Discovery Rate^32^.

### Golgi staining and analysis

Brains from 3-month-old mice were stained using the FD Rapid GolgiStain Kit (FD Neurotechnologies, Inc). They were then flash frozen and sectioned on a cryostat at 100µm thickness and further processed as per kit instructions. Hippocampal CA1 pyramidal neurons were imaged on a laser scanning confocal microscope (Leica SP5). *z*-Stacks were obtained of whole neurons (10-15 *z-*intervals per neuron). 65 Ctrl and 51 *Atrx*-cKO neurons were traced from 3 Ctrl/*Atrx*-cKO pairs. Dendrites were analyzed in ImageJ (FIJI) using the Simple Neurite Tracer plugin; the traces were analyzed using the Sholl plugin in ImageJ (FIJI) at a radius step size of 4µm^33,34^. Statistics were calculated by two-way repeated measures ANOVA with Sidak’s multiple comparison test or unpaired Student’s T-tests where applicable.

### Open field test

In the open field test, locomotor activity was automatically recorded (AccuScan Instrument). The mice were placed in an arena with an area of 20 cm × 20 cm with 30 cm high walls. Mice were acclimated to the room for ten minutes prior to testing. Locomotor activity was measured in 5 min intervals over 2 hours as previously described^17^. Distance travelled and time spent in the center were recorded. Statistics were calculated by two-way repeated measures ANOVA with Sidak’s multiple comparison test or unpaired Student’s T-tests where applicable.

### Elevated plus maze

Animals were placed in the center of the elevated plus maze (Med Associate Inc) and their activity was recorded over 5 min. Total time spent in the open and closed arms was recorded using computer software (AnyMaze). The center of the mouse body was used as an indicator of which zone they were in. Statistics were calculated by unpaired Student’s T-tests.

### Y maze

Animals were placed in the center of a symmetrical three-armed Y maze as described^17,35^. Each mouse underwent one trial of 5 minutes. Order of arm entry was recorded using computer software (AnyMaze) and spontaneous alternation was counted when a mouse entered all three arms in a row without visiting a previous arm.

### Novel object recognition

Mice were habituated in an open arena with no objects for 5 minutes on Day 1 and Day 2, as described^17^. On Day 3, mice were exposed to two identical objects for ten minutes (A; a red plastic ball attached on top of a yellow cube base). Video tracking (AnyMaze) was used to record time spent with each object. To test short-term memory, mice were exposed to the original object (A) and a novel object (B; a blue plastic pyramid attached on top of a green prism base) 1.5 hours after training. To test long-term memory, mice were exposed to (A) and (B) 24 hours after training. Novel object recognition was expressed as the percentage of time spent with the novel object as a fraction of the total time spent interacting. Interaction was defined as sniffing or touching the object, but not leaning against or climbing on the object.

### Morris water maze

The Morris water maze was conducted as described previously^36^. The task was performed in a 1.5 m diameter pool with 25°C water and the platform was submerged 1 cm beneath the water surface. Spatial cues (shapes printed in black & white) were distributed around the pool. Mice were given four trials (90 s) a day for 4 consecutive days with a 15 min intertrial period. The latency, distance, and swim speed to find the platform was recorded using video tracking software (AnyMaze). If the mice did not find the platform within 90 s, they were gently guided onto the platform. On the fifth and the twelfth days the platform was removed and time spent in each quadrant of the maze was recorded using the video software. Statistics were calculated by one- or two-way repeated measures ANOVA with Sidak’s multiple comparison test, where applicable.

### Contextual fear conditioning

To measure contextual fear memory, mice were placed in a 20 cm × 10 cm clear acrylic enclosure with a metal grid floor and one wall distinct from the others (stripes were drawn on one of the walls). The chamber was equipped with an electric shock generator. Videos were recorded using the AnyMaze video tracking software. On Day 1, mice could explore the enclosure freely and at 150 s the mice were given a shock (2 mA, 180 V, 2 s). After 30 s, the mice were returned to their home cage. The next day (24 h), the mice were placed back into the enclosure for 6 min and freezing time was measured in 30 s intervals. Freezing was defined as immobility lasting more than 0.5 s. Statistics were calculated by two-way repeated measures ANOVA with Sidak’s multiple comparison test or unpaired Student’s T-tests where applicable.

### Touchscreen assays

The paired associate learning (dPAL) and visual paired discrimination (VPD) and reversal tasks were performed as previously described^37-39^. Animals were food-restricted to 85% starting body weight. Animals were separated into two counter-balanced subgroups to control for time of day of testing, and equipment variation. Mice were tested in Bussey-Saksida mouse touch screen chambers (Lafayette Neuroscience) with strawberry milkshake given as a reward.

For the dPAL acquisition phase, animals were tested for their ability to associate objects with locations. Mice were presented with two images in two of three windows; one image was in its correct location (S+) and one was in one of its two incorrect locations (S-). The third window was blank. A correct response triggered reward presentation and start of an inter-trial period. The pre-training was repeated until mice reached criterion (completion of 36 trials within 60 minutes). The dPAL evaluation phase was performed for 45 sessions over 9 weeks. A correct response triggered reward presentation, whereas an incorrect response caused a 5 s time out and the house lights to turn on. An incorrect response also resulted in a correction trial, where the same S+/S-images were presented in the same two locations until the mouse responded correctly. The mouse was given 36 trials over 60 minutes per day. Percent correct, number of correction trials, latency to a correct or incorrect response, and latency to retrieve reward were recorded for each week.

VPD acquisition required the animal to touch the same image (S+) no matter which window it appeared in. The other screen had an incorrect image (S-). A correct response triggered reward presentation, whereas an incorrect response triggered the house lights to turn on, a time out of 5 s, and a correction trial to begin (previous trial repeated until a correct choice is made). This was repeated until mice reached criterion of 24/30 trials correct within 60 minutes over 2 consecutive days, after which baseline measurements were done for two sessions. Parameters for baseline were identical to the acquisition steps.

Immediately following baseline measurements, the VPD task reversal began, where most parameters were the same as the acquisition, but the correct image associated with the reward was S-, and the incorrect response that triggers house lights was S+. The mouse was given 30 trials per day over 10 days. Percent correct, number of correction trials needed, latency to a correct or incorrect response, and latency to retrieve reward on each day were recorded. Statistics were calculated by two-way repeated measures ANOVA with Sidak’s multiple comparison test or unpaired Student’s T-tests where applicable.

### RNA sequencing

Hippocampal total RNA was isolated using the miRVANA RNA isolation kit (ThermoFisher) from three pairs of 3-month-old control and *Atrx*-cKO male and female mice (12 samples total). RNA was quantified using the Qubit 2.0 Fluorometer (Thermo Fisher Scientific, Waltham, MA) and quality was assessed using the Agilent 2100 Bioanalyzer (Agilent Technologies Inc., Palo Alto, CA) with the RNA 6000 Nano kit (Caliper Life Sciences, Mountain View, CA). Libraries were prepared, including rRNA reduction, using the ScriptSeq Complete Gold Kit (H/M/R) (Illumina Inc., San Diego, CA). Samples were fragmented, cDNA was synthesized, tagged, cleaned-up and subjected to PCR with barcoded reverse primers (ScriptSeq Index PCR Primers) to permit equimolar pooling of samples into one library. The pooled library size distribution was assessed on an Agilent High Sensitivity DNA Bioanalyzer chip and quantitated using the Qubit 2.0 Fluorometer. All samples were sequenced at the London Regional Genomics Centre (Robarts Research Institute, London, Ontario, Canada; http://www.lrgc.ca) using the Illumina NextSeq 500 (Illumina Inc., San Diego, CA). The libraries were sequenced as a paired end run, 2 × 76 bp, using a High Output v2 kit (150 cycles). Fastq data files were downloaded from BaseSpace. At least 60 million fragments were obtained for each sample. Raw reads were pre-processed with the sequence-grooming tool cutadapt^40^ version 0.4.1 with the following quality trimming and filtering parameters (‘--phred33 -- length36 -q 5 --stringency 1 -e 0.1’). Each set of paired-end reads was mapped against the Mus Musculus GRCm38.p6 primary assembly downloaded from Ensembl^41^ release 94 (https://useast.ensembl.org/Mus_musculus/Info/Annotation) using HISAT2 version 2.0.4. SAMtools was then used to sort and convert SAM files. The read alignments and Mus Musculus GRCm38 genome annotation were provided as input into StringTie v1.3.3^42^ which returned gene and transcript abundances for each sample. We imported coverage and abundances for transcripts into R using the tximport^43^ R package and conducted differential analysis of transcript count data stratified or not on sex using the DESeq2 R package. For the unstratified analysis, an interaction term was added to the model in order to test if the effect of ATRX deletion differs across sex. We use the independent hypothesis weighting (IHW) Bioconductor package^44^ to weight p-values and adjust for multiple testing using the procedure of Benjamini Hochberg (BH)^45^. Functional enrichment of significant genes with aggregated p-value <0.05 was evaluated by the ‘weight01’ algorithm implemented in the R topGO package^46^ that assesses, prunes and weighs for local dependencies of gene ontology (GO) terms, and Fisher exact test.

### Transmission electron microscopy

Mice were perfused with 4% paraformaldehyde (VWR) dissolved in phosphate buffer and sectioned at 500µm on a Vibratome Series 1000 Sectioning System. Sections were post-fixed overnight in 4% paraformaldehyde for 24h and in 1% glutaraldehyde for 1 hour, then washed and left in phosphate buffer until preparation of ultra-thin sections. Slices were transferred to the Biotron facility at Western University for the remaining steps. Coronal slices from the hippocampal layer were rinsed in distilled water, post-fixed for one hour at room temperature in 1% osmium tetroxide (Electron Microscopy Sciences Warrington, PA) and 1.5% potassium ferrocyanide and post-fixed for a second hour in 1% osmium tetroxide^47^. The slices were quickly dehydrated through a graded series of ethanol, rinsed in 100% acetone and then infiltrated with inversion for one hour in 50% epon-araldite resin and placed overnight in 100% resin on a rotator. The slices were polymerized overnight at 60°C in a vented oven, sandwiched between two sheets of Aclar film covered with a light weight^48^. The hippocampal region was cut from the resin-embedded slice using a razor blade and mounted on a blank epoxy block using cyanoacrylate glue. Semi-thin sections were made with a Reichert Ultracut ultramicrotome and stained with toluidine blue and used to select the CA1 neuropil region for ultrathin sectioning. Thin sections (100nm) were collected on 200 mesh nickel grids (EMS), post-stained with lead citrate and images were collected with a Philips 420 transmission electron microscope equipped with an AMT 4K megapixel XR41S-B camera (Woburn, MA). Images were captured within the stratum radiatum / stratum lacunosum moleculare at 9200X in a semi-random manner to obtain at least 50 synapses containing a distinct synaptic cleft without regard to the length of the cleft. Image analysis was performed in ImageJ. A total of 104 Ctrl and 84 *Atrx-*cKO synapses were quantified from 3 Ctrl/*Atrx*-cKO pairs. Synapses were binned in 50nm increments from the active zone and the number of docked vesicles and vesicles in each bin were counted; this also determined total number of vesicles per synapse. Vesicle cluster size was measured to calculate vesicle density. The area of the post-synaptic density was also quantified. Statistics were calculated by two-way repeated measures ANOVA with Sidak’s multiple comparison test or unpaired Student’s T-tests where applicable.

### Statistical analyses

All data were analyzed using GraphPad Prism software with Student’s T test (unpaired, two-tailed) or one or two-way repeated measures ANOVA with Sidak’s post-test where applicable. All results are depicted as mean +/− SEM unless indicated otherwise. P values of less than 0.05 were considered to indicate significance.

## Results

### *Generation and validation of mice with neuron-specific* Atrx *deletion*

We generated mice lacking ATRX in postnatal forebrain excitatory neurons by Cre/loxP mediated recombination of the mouse *Atrx* gene with the *CaMKII-*Cre driver line of mice^49^. To confirm loss of ATRX, we performed immunofluorescence staining of control and conditional knockout (*Atrx*-cKO) brain cryosections obtained from 3-month-old mice (**Figure 1a,b**). ATRX is highly expressed in excitatory neurons of the hippocampus of control mice, including the cortex and hippocampal CA1, CA2/3, and dentate gyrus neurons, but is absent in these cells in the *Atrx*-cKO mice. Additional validation of *Atrx* inactivation in *Atrx*-cKO mice was achieved by qRT-PCR (**Figure 1c**), showing that *Atrx* expression is decreased by 78% (+/− 9.4%) and 81% (+/− 1.7%) in the cortex and the hippocampus, respectively, which is expected from a neuron-specific deletion. The brain sub-region specificity of ATRX loss was demonstrated by western blot analysis, showing reduced protein levels in the rostral and caudal cortices and hippocampus, but not in the cerebellum (**Figure 1d**). The mice survived to adulthood and had normal general appearance and behaviour. However, body weight measurements revealed a small but significant reduction in *Atrx*-cKO compared to control mice (**Figure 1e**). These findings demonstrate that we achieved specific deletion of *Atrx* in excitatory neurons and while the mice were slightly smaller, they survived to adulthood, allowing further analyses in the adult brain.

**Figure 1:**
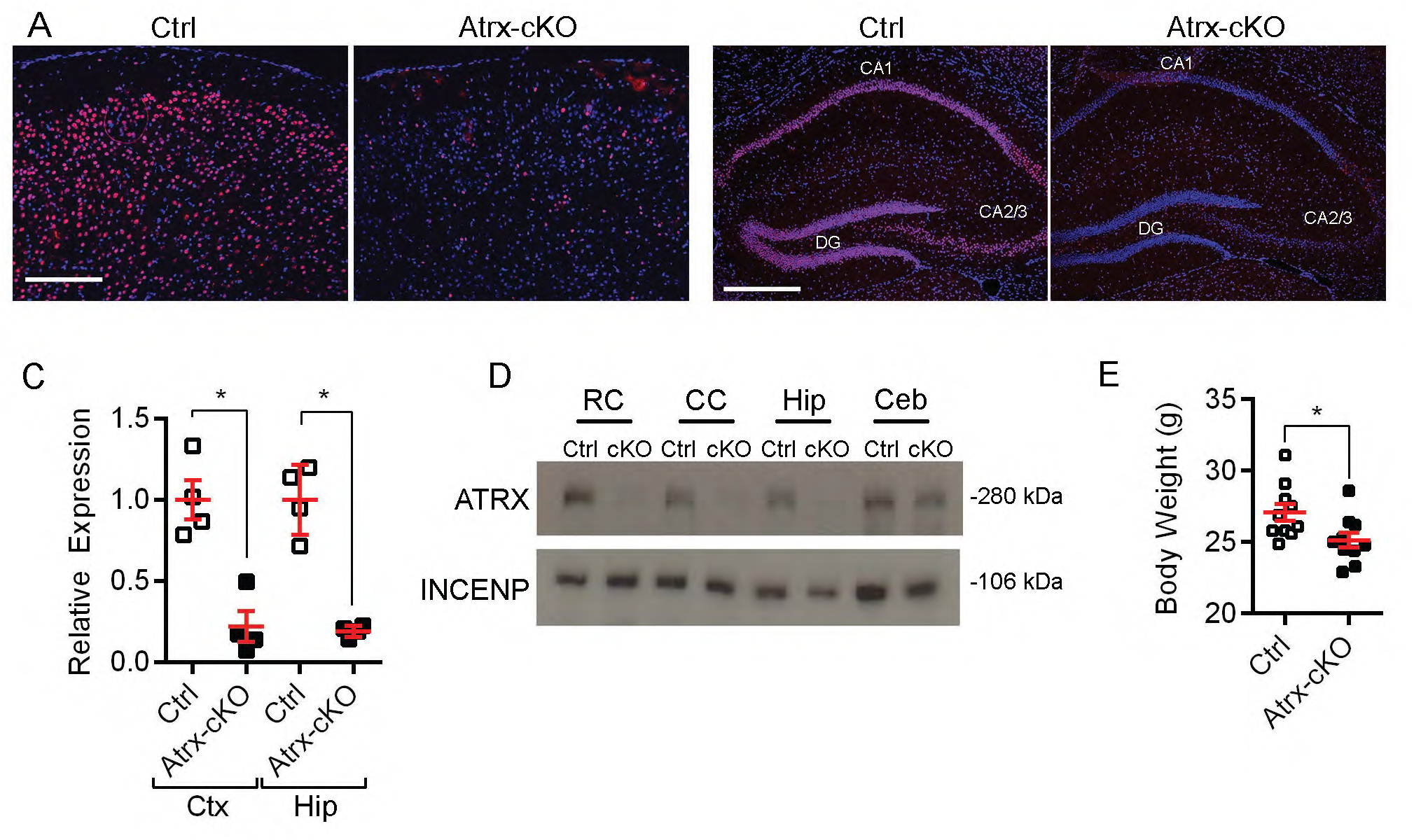
Validation of *Atrx* inactivation in pyramidal neurons of the forebrain. (A) Immunofluorescence in cortex of control (Ctrl) and knockout (*Atrx*-cKO) male mice. ATRX: red; DAPI: blue. Scale bar: 100 µm. (B) Immunofluorescence in hippocampus of control (Ctrl) and knockout (*Atrx*-cKO) male mice. ATRX: red; DAPI: blue. Scale bar: 200 µm. (C) *Atrx* RNA transcripts measured by qRT-PCR in the rostral cortex (Ctx) (p<0.005) and hippocampus (HI) (p<0.001) (n=4). (D) Western blot of ATRX and INCENP using protein extracts from rostral cortex (RC), caudal cortex (CC), hippocampus (Hip) and cerebellum (Ceb). (E) Body weight of control and *Atrx*-cKO male mice at 3 months of age (n=10) (p<0.05). Statistics by unpaired Student’s T-test.

### MRI analysis reveals *anatomical abnormalities in the hippocampus of Atrx-cKO mice*

We first examined control and *Atrx*-cKO mouse brains for neuroanatomical anomalies by magnetic resonance imaging (MRI). Using a T2-weighted MRI sequence, we were able to analyze and compare the entire brain as well as independent brain regions from 16 control and 13 Atrx-cKO male animals. The data obtained showed that the overall volume of the *Atrx*-cKO brain is significantly smaller compared to controls (92.8% of control volume, P<0.0001), as indicated by whole volume in mm^3^ and cumulative serial slices of control and *Atrx-*cKO brains (**Figure 2a,b**), which correlates with the smaller body size of the mice. Due to the reduction in body size and absolute total brain and hippocampal volumes of the *Atrx*-cKO mice, we next examined hippocampal neuroanatomy relative to total brain volume (**Figure 2c**). The relative volume of the CA1 stratum radiatum (SR) and stratum lacunosum moleculare (SLM) was significantly increased in the *Atrx-*cKO mice compared to controls (**Figure 2d**) whereas the CA3 pyramidal layer was significantly decreased in size (**Figure 2e**). Relative volumes of all hippocampal regions are tabulated in **Figure 2f**.

**Figure 2:**
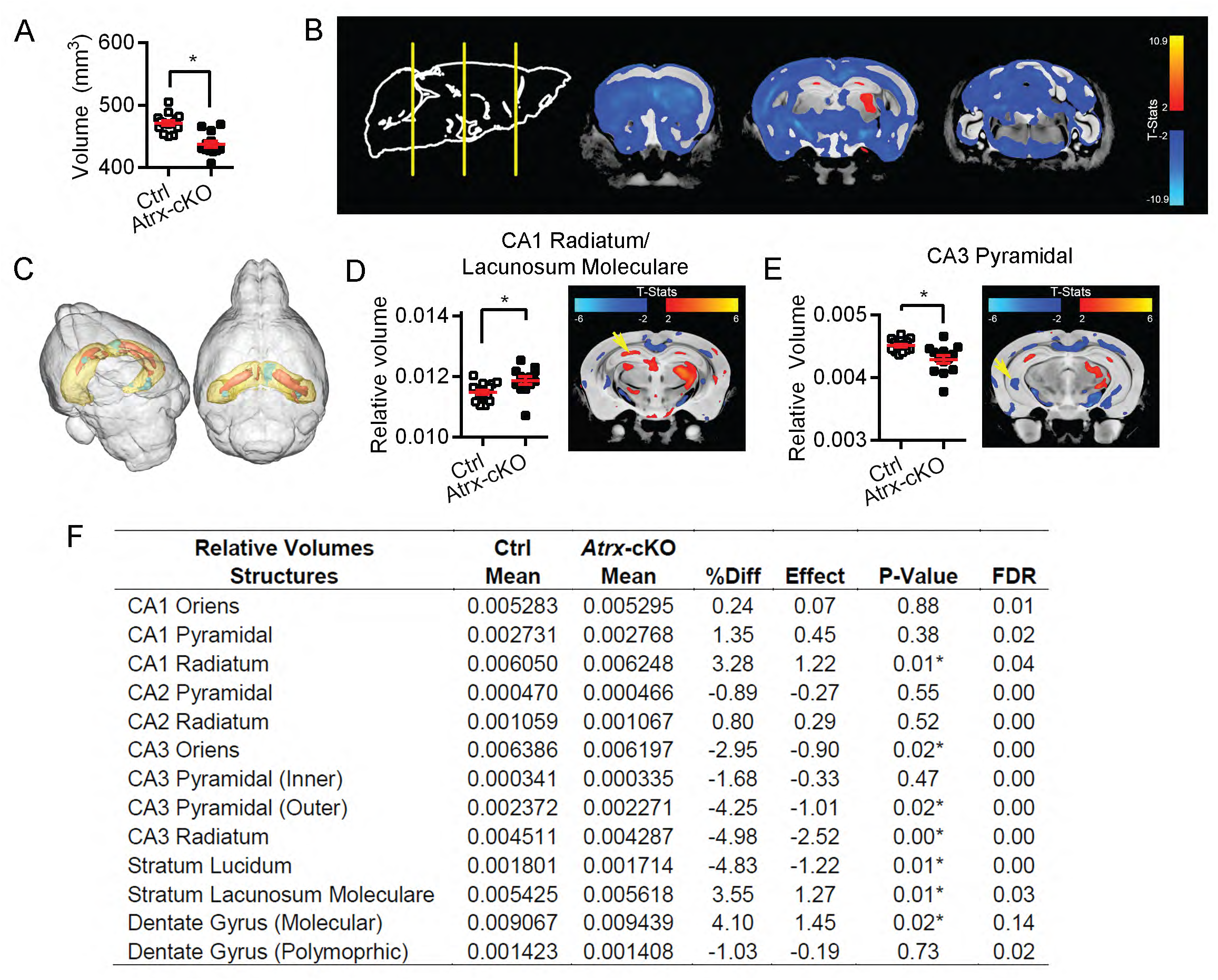
MRI reveals altered morphology of *Atrx*-cKO hippocampi. (A) Absolute volumes of Ctrl and *Atrx*-cKO mouse brains in mm3 (p<0.0001). (B) Cumulative images of Ctrl and *Atrx*-cKO brains displaying changes in density of absolute volume. (C) Cumulative 3D image from Ctrl and *Atrx*-cKO mouse brains displaying changes in density of hippocampus volume relative to brain size. Hippocampus is colored yellow, areas of increased volume in *Atrx*-KO are colored orange and areas of decreased volume in green. Relative size of CA1 stratum radiatum and lacunosum moleculare in Ctrl and *Atrx*- cKO mouse brains (p<0.05), with MRI image showing cumulative changes in *Atrx*-cKO compared to control. (E) Relative size of CA3 pyramidal layer in Ctrl and *Atrx*-cKO mouse brains (p<0.005), with MRI image showing cumulative changes in *Atrx*-cKO compared to control. (F) Relative volumes of all hippocampal structures, including mean and standard deviation (SD) of Ctrl and *Atrx*-cKO, percent difference (%Diff), effect size, P-value, and false discovery rate (FDR). Asterisks indicate p<0.05. All data is derived from 16 Ctrl and 13 *Atrx*-cKO mouse brains.

We postulated that the increase in relative volume of the CA1 SR/SLM may be due to increased length or branching of CA1 apical dendrites. To investigate this possibility, Golgi staining was used to sporadically label neurons (**Figure 3a**) and Sholl analysis was performed on confocal microscopy images to evaluate apical dendrite branching of CA1 hippocampal neurons. However, no significant difference in dendritic branching or length was observed between control and *Atrx*-cKO mice, whether analyzed separately for apical or basal dendrites (**Figure 3b-g**). Increased relative volume might also be caused by an increased number of cells, but immunofluorescence staining and quantification of astrocytes (GFAP+) and microglia (IBA1+) and total number of cells (inferred from DAPI+ staining) revealed no differences in *Atrx*-cKO hippocampi (**Figure 3h-j**). Overall, the increased relative volume of the CA1 SR/SLM cannot be explained by increased length or complexity of dendritic trees or by an increased number of glial cells.

**Figure 3:**
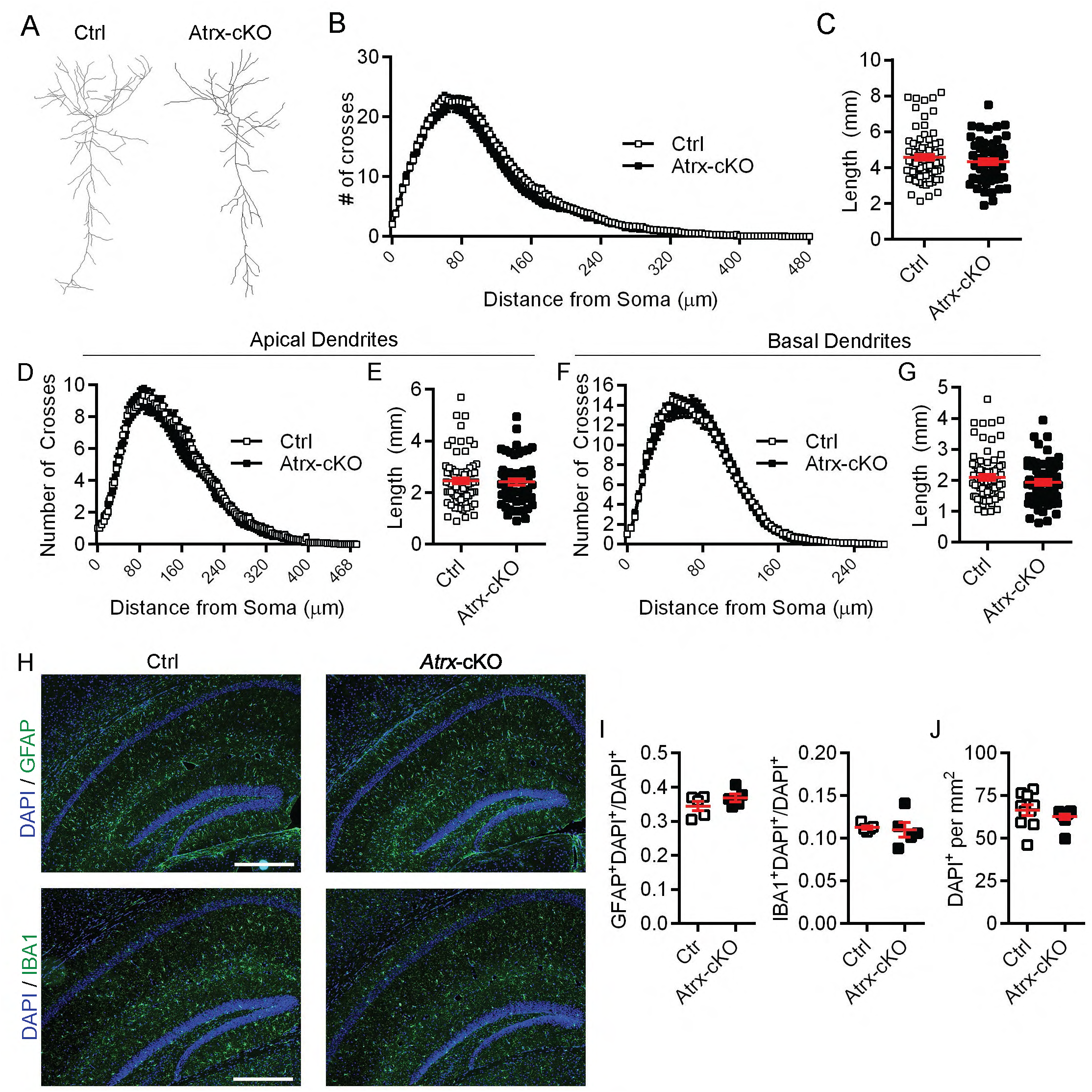
CA1 dendritic length and branching and the number of non-neuronal cells are not affected in *Atrx*-cKO mouse hippocampi. (A) Representative Golgi traces from Ctrl and *Atrx*-cKO hippocampal CA1 pyramidal neurons. (B) Sholl analysis of Ctrl (n=73) and *Atrx*-cKO (n=53) CA1 pyramidal neurons (p=0.0842). (C) Total length of Ctrl and *Atrx*- cKO CA1 dendrites (p=0.2994). (D) Sholl analysis of Ctrl and *Atrx*-cKO CA1 apical dendrites (p=0.3858). (E) Total length of Ctrl and *Atrx*-cKO CA1 apical dendrites (p=0.7832). (F) Sholl analysis of Ctrl and *Atrx*-cKO CA1 basal dendrites (p=0.7339). (G) Total length of Ctrl and *Atrx*-cKO CA1 basal dendrites (p=0.2237). (H) Immunofluorescence staining of GFAP or IBA1 in Ctrl and *Atrx*-cKO hippocampi. DAPI was used as a counterstain. Scale bar indicates 400 µm. (I) Quantification of the proportion of GFAP+ (p=0.2151) and IBA1+ (p=0.7903) cells in stratum radiatum/stratum lucidem moleculare of Ctrl and *Atrx*-cKO hippocampi (n=5). (J) Quantification of DAPI+ cells per mm2 in n stratum radiatum/stratum lucidem moleculare of Ctrl and *Atrx*-cKO hippocampi (n=10, p=0.2904). Data was analyzed by unpaired Student’s T-Test or two-way repeated measures ANOVA with Sidak’s multiple comparisons test where applicable, and asterisks indicate p<0.05. Data is displayed as mean +/-SEM.

### *Pre- and post-synaptic structural defects in* Atrx*-cKO male mice*

Based on the hippocampal structural alterations we detected by MRI, we looked more closely at potential ultrastructural changes in the CA1 SR/SLM area using transmission electron microscopy (TEM) (**Figure 4a**). The presynaptic boutons were divided in 50nm bins from the active zone, and the number of vesicles in each bin was counted. The spatial distribution of vesicles in relation to the cleft was unchanged between the *Atrx*-cKO mice and controls (**Figure 4b**). However, we found that the total number of vesicles, the density of the vesicles, and the number of docked vesicles was significantly decreased at *Atrx-*cKO compared to control synapses (**Figure 4c-e**). We also analysed other structural aspects of synapses and found that the size of the post-synaptic density and the width of the synaptic cleft were both increased in *Atrx-*cKO compared to controls (**Figure 4f,g**). The length of the active zone, cluster size, or diameter of the vesicles did not vary significantly between control and *Atrx*-cKO samples (**Figure 4h-j**). These results suggest that ATRX is required for structural integrity of the pre- and post-synapse, including maintenance of the synaptic vesicle pool at pre-synaptic termini and potential defects in postsynaptic protein clustering.

**Figure 4:**
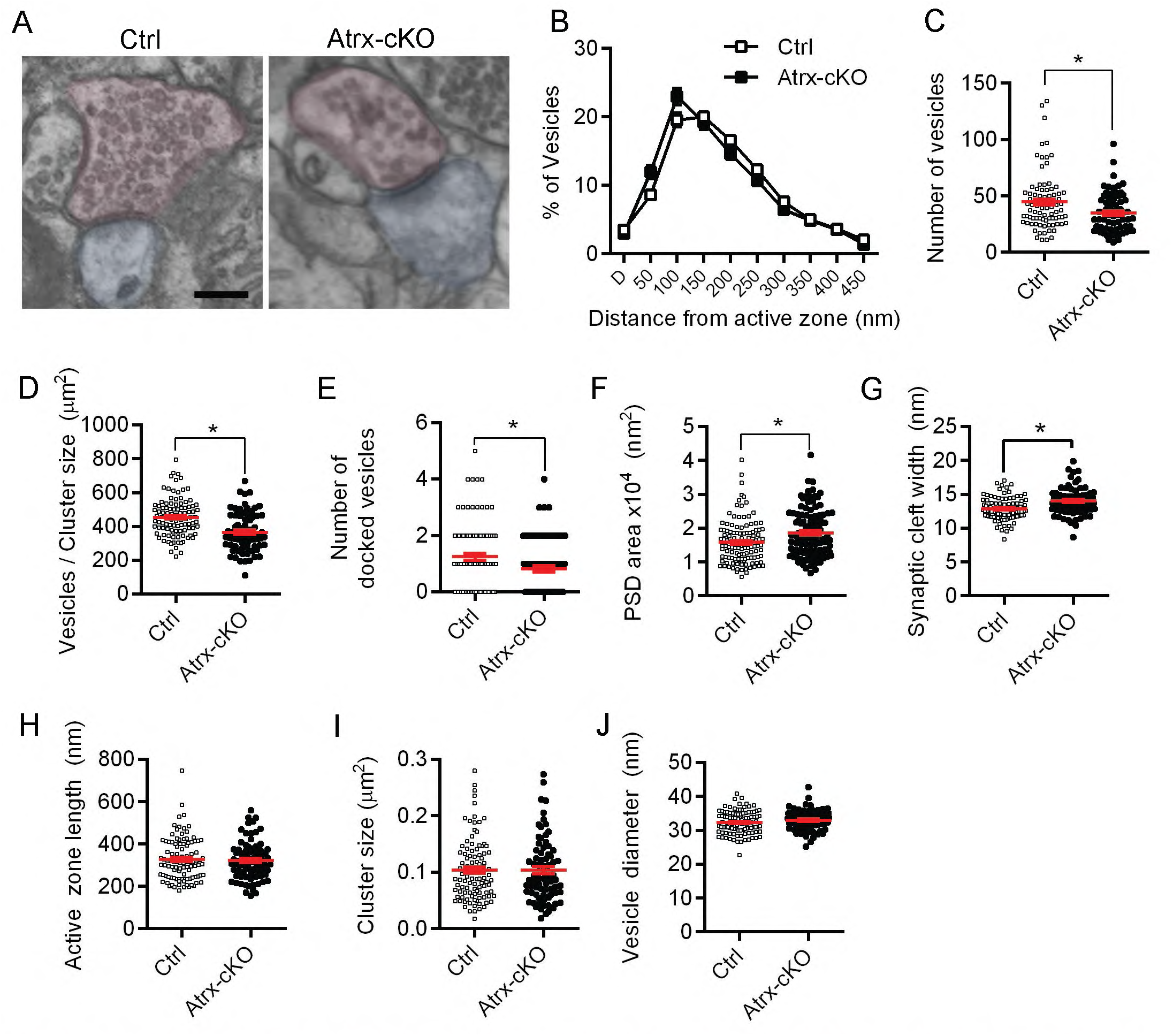
Ultrastructural analysis of *Atrx*-cKO CA1 apical synapses reveals a reduced number of total or docked presynaptic vesicles, wider synaptic cleft and larger post-synaptic density. (A) Representative images of Ctrl and *Atrx*-cKO CA1 synapses imaged by transmission electron microscopy. Red shading indicates presynaptic neuron, blue shading indicates postsynaptic neuron. Scale bar = 200 nm. (B) Number of vesicles in 50nm bins from the active zone (p=0.9504) (Ctrl n=60, *Atrx*-cKO n=63). (C) Number of total vesicles (p<0.01). (D) Density of vesicles per cluster (p<0001). (E) Number of docked vesicles (p<0.05). (F) Post-synaptic density (PSD) area (p<0.005). (G) Synaptic cleft width (p< 0.0001). (H) Length of the pre-synaptic active zone (p=0.7039). (I) Synaptic vesicle cluster size (p=0.9917). (J) Vesicle diameter (p=0.1895). Data was analyzed by unpaired Student’s T-test or two-way repeated measures ANOVA with Sidak’s multiple comparisons test where applicable, and asterisks indicate p<0.05. Data is displayed as mean +/− SEM.

### Loss of ATRX in neurons leads to long-term spatial learning and memory deficits

We next performed a battery of behaviour tests on male *Atrx*-cKO mice. Locomotor activity in the open field test was not significantly different between *Atrx*-cKO mice compared to controls, either over time (F=0.1722, P=0.6803) or if considering total distance travelled (T=0.6691, P=0.5072; **Supp. Figure 1a, b**). The *Atrx*-cKO mice did not spend significantly more time in the centre of the chamber over time (F=2.960, P=0.0927; **Supp. Figure 1c**). However, they spent significantly more total time in the centre of the chamber when compared to controls (T=2.262, P=0.0291; **Supp. Figure 1d**), suggesting that loss of ATRX in neurons has an anxiolytic effect. This was confirmed by the behaviour of the *Atrx*-cKO mice in the elevated plus maze, as they spent significantly more time in the open arm of the maze compared to controls (T=2.158, P=0.0403; **Supp. Figure 1e,f**). We observed no difference in percent alternation in the Y-maze (T=0.9431, P=0.3543; **Supp. Figure 1g**), nor in the training phase of the novel object recognition task (Ctrl T=1.128, P=0.2697; cKO T=1.657, P=0.1096; **Supp. Figure 1h**) or memory tests at 1.5 hours (Ctrl T=4.277, P<0.001; cKO T=3.545, P<0.005) or 24 hours (Ctrl T=1.645, P=0.112; cKO T=2.459, P<0.05; **Supp. Figure 1i**).

To investigate the effects of neuronal-specific ATRX ablation on spatial learning and memory, we tested the mice in the Morris water maze task. The *Atrx*-cKO mice showed a significant delay in latency to find the platform on day 3 of the learning portion of the task; however, by day 4 they were able to find the platform as quickly as the control mice (F=4.622, P=0.0404; **Figure 5a**). This finding was reflected in the distance traveled to find the platform (F=4.829, P=0.0364; **Figure 5b**). Swim speed was comparable to controls over the four days of learning the task (F=0.04238, P=0.8384; **Figure 5c**), confirming the findings of the open field test which showed similar activity levels between control and *Atrx*-cKO mice. Memory was tested on day 5 (24h after last training day) and day 12 (7 days after last training day). Both controls (F=29.36, P<0.0001) and the *Atrx*- cKO male mice (F=18.97, P<0.0001) spent significantly more time in the target quadrant than the left, opposite, or right quadrants (**Figure 5d**). However, on day 12, *Atrx*-cKO male mice failed to spend significantly more time in the target quadrant (F=1.420, P=0.2594) whereas controls were still able to remember (F=6.785, P<0.01), suggesting a long-term spatial memory deficit (**Figure 5e**). Moreover, *Atrx*-cKO male mice froze significantly less in the contextual fear memory task in comparison to their control counterparts 24h after a foot shock (over time F=5.392, P=0.0251; total time T=2.322; P=0.0251; **Figure 5f, g**). These behavioural analyses suggest that ATRX in required in excitatory neurons for long-term hippocampal-dependent spatial learning and memory.

**Figure 5:**
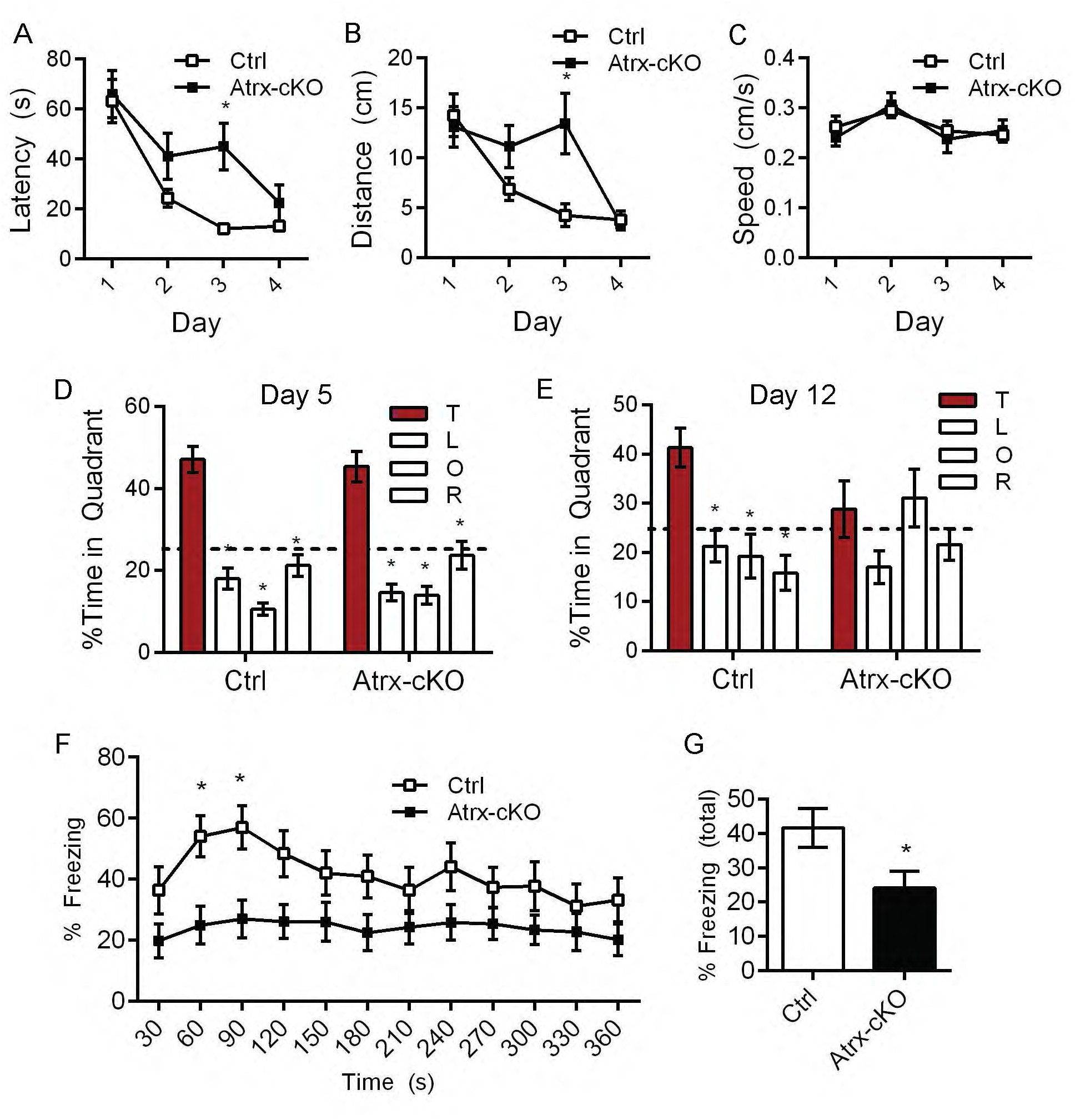
The *Atrx*-cKO mice exhibit impaired long-term spatial memory in the Morris water maze paradigm and in the contextual fear conditioning task. (A,B,C) Latency to reach the platform (p<0.05), distance travelled (p<0.05), and swimming speed (p=0.8384) over four days (four trials per day) in the Morris water maze (n=15). (D) Percent time spent in each quadrant after removal of the platform on day 5. Dotted line indicates chance at 25%. (E) Percent time spent in each quadrant after removal of the platform on day 12. Dotted line indicates chance at 25%. (F) Percent of time freezing over 360s during the contextual fear conditioning task (p<0.05) (n=22). (G) Total time freezing (p<0.05). Statistics by two-way repeated measures ANOVA with Sidak’s multiple comparisons test or Student’s T-test where applicable, asterisks indicate p<0.05.

### Neuron-specific deletion of Atrx in female mice does not cause memory deficits

To determine whether loss of ATRX in female mice would exhibit similar behavioural defects as seen in male mice, we generated *Atrx*-cKO female mice (*Atrx*-cKO^Fem^) and validated loss of ATRX in forebrain excitatory neurons by immunofluorescence staining and qRT-PCR (**Supplementary Figure 2a-c**). *Atrx*-cKO^Fem^ mice displayed normal locomotion over time (F=1.239, P=0.2796) or in total distance travelled in the open field test (T=1.113, P=0.2796). Furthermore, we observed no changes in anxiety in the open field test over time (F=0.009, P=0.9254), total time spent in centre (T=0.095, P=0.9254; **Supp. Figure 3a-d**) nor in the time spent in the open arm (T=0.4947, P=0.6250) and closed arm (T=0.4907, P=0.6277; **Supp. Figure 3e**) in the elevated plus maze in the *Atrx*-cKO^Fem^ mice compared to control, indicating that the decreased anxiety levels are sex-specific.

Conversely to what was observed in the *Atrx*-cKO male mice we failed to observe differences between *Atrx*-cKO^Fem^ and control mice in latency in the Morris water maze training sessions (latency F=3.631, P=0.0683; distance F=1.385, P=0.2503; speed F=1.243, P=0.2754; **Supp. Figure 4a-c**). Memory of the platform location remained equivalent to that of controls when tested on day 5 or day 12 (Ctrl day 5 F=12.80, P<0.0001; cKO day 5 F=11.63, P=0.0006; Ctrl day 12 F=16.47, P<0.0001; *Atrx*-cKO^Fem^ day 12 F=12.97, P<0.0001; **Supp. Figure 4d, e**). Finally, there was no difference in freezing between control and *Atrx*-cKO^Fem^ 24h after foot shock in the contextual fear conditioning task (over time F=0.02257, P=0.8818; total time T=0.1502, P=0.8818; **Supp. Figure 4f, g**). We conclude that learning and memory is not impaired by ATRX loss in forebrain excitatory neurons of the *Atrx*-cKO^Fem^ mice.

### Impaired object location associative memory in the rodent version of the paired associate learning (dPAL) task

Given the observed male-specific defects in spatial memory, we performed additional translational cognitive tasks on the *Atrx*-cKO male mice. The dPAL touchscreen task in mice is analogous to cognitive testing done in humans by the Cambridge Neuropsychological Test Automated Battery (CANTAB)^50,51^ and normal performance in this task is thought to partly depend on the hippocampus^39,52^.

Control and *Atrx*-cKO mice were trained to identify the position of three images as depicted in **Figure 6a**, undergoing 36 trials per day for 10 weeks. The results demonstrate that the *Atrx*-cKO mice exhibit a profound deficit in this task, indicated by both the percent correct (F=10.53, P=0.0031; **Figure 6b**) and the number of correction trials required (F=30.64, P<0.0001; **Figure 6c**) (**Supplementary video 1**, **2**). These defects were not due to an inability to perform within the chamber or to attentional deficits, as latency to a correct answer (F=0.4802, P=0.4943), to an incorrect answer (F=0.1259, P=0.7255), and to retrieve the reward (F=0.9840, P=0.3300) was not significantly different between control and *Atrx*-cKO mice (**Figure 6d-f**). To determine if the impairment in the dPAL task is caused by a vision problem rather than a learning defect, the mice were also tested in the visual paired discrimination (VPD) touchscreen task which requires the mice to discriminate between two images regardless of position on the screen. While the *Atrx*- cKO mice took significantly longer to reach the criterion pre-testing (T=2.945, P=0.0067; **Figure 6g**), there was no difference in the percent correct during baseline tests or after reversal compared to controls, suggesting that vision is intact in the *Atrx*-cKO mice (F=1.388, P=0.2490; **Figure 6h**). They did however require an increased number of correction trials, indicating that their cognitive flexibility may be marginally impaired compared to controls (F=11.84, P=0.0019; **Figure 6i**). The results of these touchscreen tests reinforce the findings that the *Atrx-*cKO male mice have impaired learning and memory.

**Figure 6:**
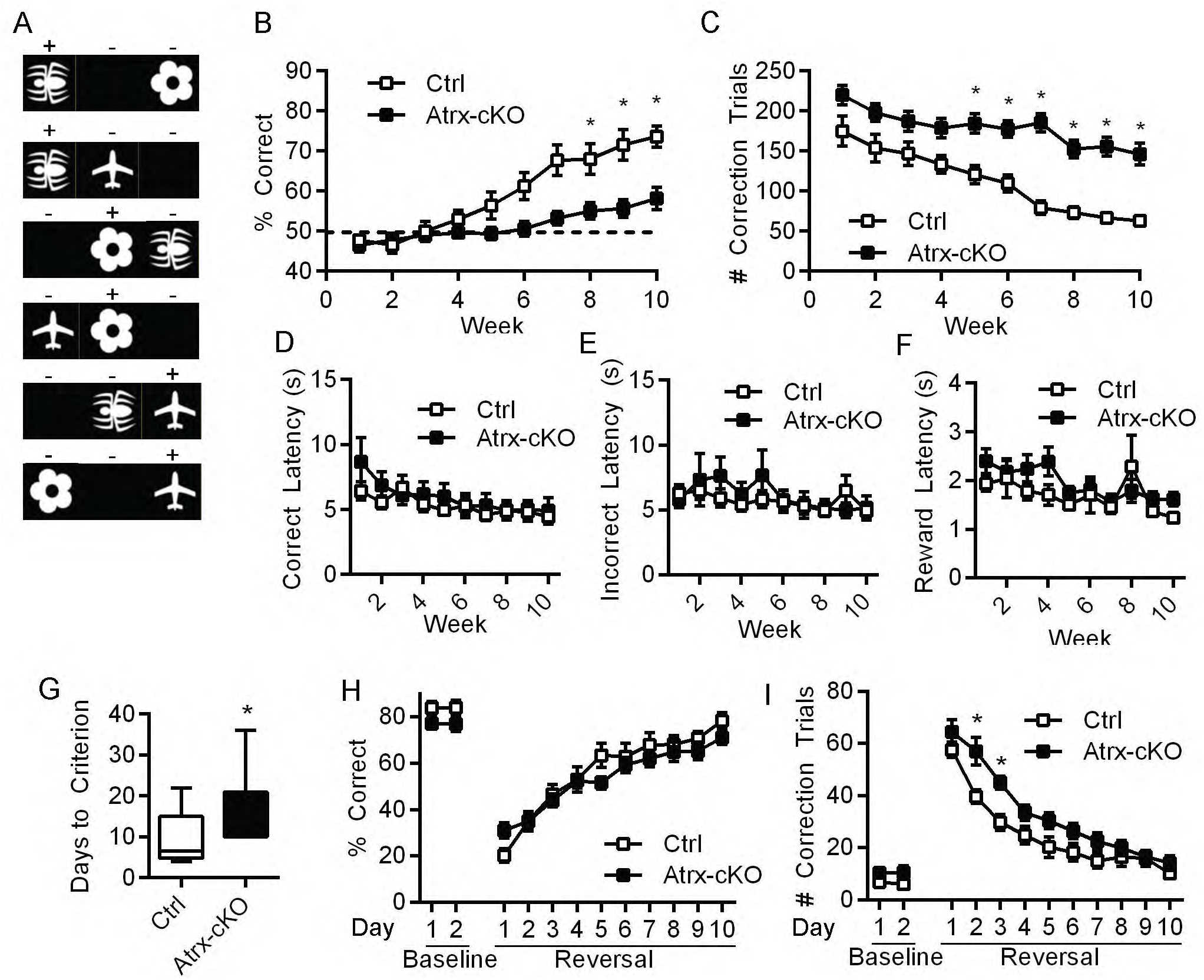
*Atrx*-cKO mice display deficits in spatial learning in the Paired-Associate Learning operant task. (A) Representative images used in the paired-associate learning task, where touching the (+) stimuli on the screen results in reward and the (-) stimuli results in negative reinforcement (Talpos et al., 2009). (B) Percent correct over 10 weeks (p<0.005). Dotted line indicates % correct by chance. (C) Number of correction trials required over ten weeks (p<0.0001). (D) Latency to choose a correct answer (p=0.49). (E) Latency to choose an incorrect answer (p=0.72). (F) Latency to retrieve the reward (p=0.33). (G) Number of days to reach criterion in the Visual Paired Discrimination task (p<0.01). (H) Percent correct during baseline (two days) and reversal (10 days) (p=0.25). (I) Number of correction trials required during baseline and reversal trials (p<0.005). Statistics by two-way repeated measures ANOVA with Sidak’s multiple comparisons test or Student’s T-test where applicable, asterisks indicate p<0.05.

### RNA sequencing of the hippocampus reveals sex-specific transcriptional changes

To identify the molecular mechanism(s) leading to spatial memory impairment, we performed RNA-sequencing in both male and female hippocampi obtained from three pairs of littermate-matched Ctrl/*Atrx-*cKO and Ctrl^Fem^/*Atrx*-cKO^Fem^ mice. There were 1520 transcripts differentially expressed in the *Atrx-cKO* males compared to control counterparts and 9068 transcripts in *Atrx-cKO*^Fem^ compared to the female controls (FDR < 0.20). To isolate transcripts that were likely to be causative to the impaired learning and memory phenotype in the male mice which was not found in the female mice, we focused on transcripts whose changes in expression with the *Atrx-cKO* were differential between male and female mice (n = 1054 transcripts, interaction term FDR < 0.05). The expression heat map of these transcripts illustrates that their expression levels are similar in control males and females but are differentially expressed when ATRX is lost depending on sex (Figure 7a). We then utilized PANTHER^53^, a tool for gene enrichment analysis based on functional annotations to examine Gene Ontology biological processes for which our list of transcripts was enriched (Figure 7b). The top five pathways included neurotransmitter receptor transport to postsynaptic membrane, protein localization to postsynaptic membrane, non-motile cilium assembly, and vesicle-mediated transport to the membrane. Therefore, the RNA sequencing revealed many transcripts related to synapses, supporting the TEM data.

**Figure 7:**
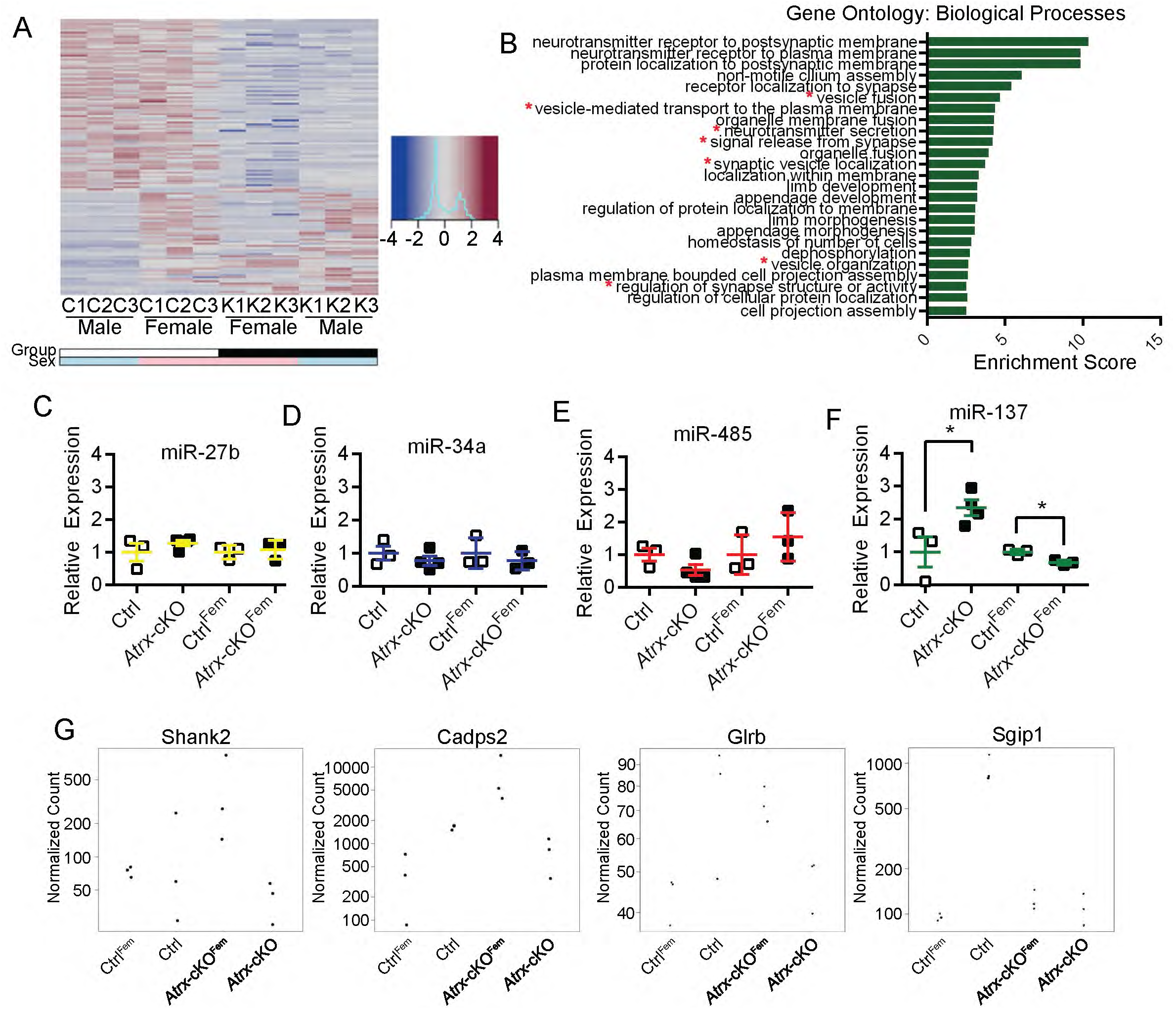
Transcriptional profiling reveals dysregulation of presynaptic vesicular genes possibly resulting from miR-137 overexpression. (A) Heat map analysis of differentially expressed transcripts according to sex (FDR < 0.05) by RNA sequencing from Ctrl, *Atrx*-cKO, Ctrl^Fem^, and *Atrx*-cKO^Fem^ hippocampi. (B) Unique transcripts that were regulated in a sex-specific manner upon loss of ATRX were used for Gene Ontology analysis and top 25 Biological Processes were listed by Enrichment value (p<0.001, FDR<0.05). Those related to synapses were noted with a red asterisk. (C) Expression of miR-485 in Ctrl and *Atrx*-cKO (p=0.1284), Ctrl^Fem^ and *Atrx*-cKO^Fem^ (p=0.3787) hippocampi normalized to miR-191. (D) Expression of miR-34a in Ctrl and *Atrx*-cKO (p=0.3072), Ctr^lFem^ and *Atrx*-cKO^Fem^ (p=0.7193) hippocampi normalized to miR-191. (E) Expression of miR-27b in Ctrl and *Atrx*-cKO (p=0.3953), Ctrl^Fem^ and *Atrx*-cKO^Fem^ (p=0.4968) hippocampi normalized to miR- 191. (F) Expression of miR-137 in Ctrl and *Atrx*-cKO (p<0.05), Ctrl^Fem^ and *Atrx*-cKO^Fem^ (p<0.01) hippocampi normalized to miR-191. (G) Transcript expression of *Shank2, Cadsp2, Glrb*, and *Sgip1* in Ctrl, *Atrx*-cKO, Ctrl^Fem^ and *Atrx*-cKO^Fem^ hippocampi. Data was analyzed by unpaired Student’s T-test where applicable, and asterisks indicate p<0.05. Data is displayed as mean +/− SEM.

Certain miRNA are enriched within presynaptic terminals and have been implicated in neurotransmitter release by controlling expression of SNARE and other synaptic vesicle proteins^54^. These miRNA include miR-485, miR-34a, miR-137, and miR-27b. miR-485 targets a vesicular glycoprotein SV2A and overexpression results in decreased neurotransmitter release^55^, while miR-34a also targets vesicular proteins SYT1 and SYN1, affecting miniature excitatory postsynaptic currents^56^. miR-27b regulates two-thirds of the presynaptic transcriptome by silencing expression of the negative transcriptional regulator *Bmi1^57^*, and miR-137 overexpression affects synaptic vesicle localization and LTP^58^. To determine whether any of these miRNAs might be regulated by ATRX, with differential effects in male and female cKO hippocampi, we assessed their expression by qRT-PCR. This analysis revealed no significant changes in the expression the miR-485, miR-34a, and miR-27b (**Figure 7c-e**). Conversely, we observed a striking and significant sex-specific effect of ATRX loss on the expression of miR-137 in *Atrx*-cKO hippocampi, with an upregulation of expression in the male hippocampi and a downregulation in the female hippocampi (**Figure 7f**). To further confirm this finding, we mined previous microarray data of neonatal forebrain tissue of mice lacking ATRX in the forebrain (*Atrx*^fl/fl^× *FoxG1*-Cre cross)^59^. Analysis of this data through the gene enrichment analysis tool TOPPGENE^60^ revealed enrichment of downregulated genes in Gene Ontology categories including synaptic signalling and regulation of excitatory potential (**Supp. Figure 5a**) as well as an enrichment for targets of miR-137 (**Supp. Figure 5b**). We compared the list of genes downregulated in the *Atrx*- cKO male hippocampi to those predicted to be regulated by miR-137 through miRNA.org (**Supp. Table 1**). We found *Shank2, Cadps2, Glrb*, and *Sgip1* expression to be inversely related to miR-137, with expression decreased in *Atrx*-cKO and increased in *Atrx*-cKO^Fem^. Shank2 and Glrb are both postsynaptic proteins, with Shank2 (SH3 and multiple ankyrin repeat domains 2) acting as a scaffolding protein within the PSD^61^ and GlrB (Glycine receptor beta), is the beta subunit of the glycine receptor^62^. Cadps2 and Sgip1 are found at the presynaptic terminal, where Cadps2 (Ca2+-dependent activator protein for secretion 2) regulates exocytosis of vesicles^63^ and Sgip1 (Src homology 3-domain growth factor receptor-bound 2-like interacting protein 1) is involved in clathrin-mediated endocytosis^64^. This data provides additional evidence that loss of ATRX in the cortex and hippocampus of male mice leads to increased miR-137 expression and consequent downregulation of its target genes, starting at early stages of forebrain development.

## Discussion

This study presents evidence that ATRX is required in a sex-specific manner in excitatory forebrain neurons for normal spatial learning and memory (**Figure 8**). We found that loss of ATRX in these neurons resulted in impaired long-term memory in the Morris water maze, contextual fear conditioning task, and impaired learning in the dPAL touchscreen assay. Magnetic resonance imaging revealed a higher relative volume of hippocampal CA1 SR/SLM and decreased relative volume of the CA3 pyramidal cell layer. There are two hippocampal pathways implicated in spatial learning and memory, the Schaffer collateral pathway and the temporoammonic pathway. The Schaffer collateral pathway involves the CA3 axons projecting to the CA1 medial apical dendrites^65^, whereas the temporoammonic pathway initiates in layer III of the entorhinal cortex and projects to the CA1 middle apical dendritic layer^66^. We originally hypothesized that the increased volume of the CA1 SR/SLM may be due to increased branching of pyramidal neurons, particularly the apical dendrites that project to this region. However, analysis of Golgi stained CA1 neurons failed to show abnormalities in dendritic branching that would explain the MRI data. Astrocyte and microglia infiltration have been linked with defects in learning and memory in various mouse models^67-69^. Yet after examination of these cell types by immunofluorescence staining, we saw no change in either astrocyte or microglia cell number. The reason for volume increase thus remains undetermined, but might be caused by other cell types infiltrating the hippocampus such as T cells or to increased volume of the perineuronal net (changes to the extracellular matrix), which has been shown in mouse models to help regulate learning and memory^70,71^. An alternative explanation is that volumetric measurements are normal in these areas but decreased elsewhere in the hippocampus.

**Figure 8:**
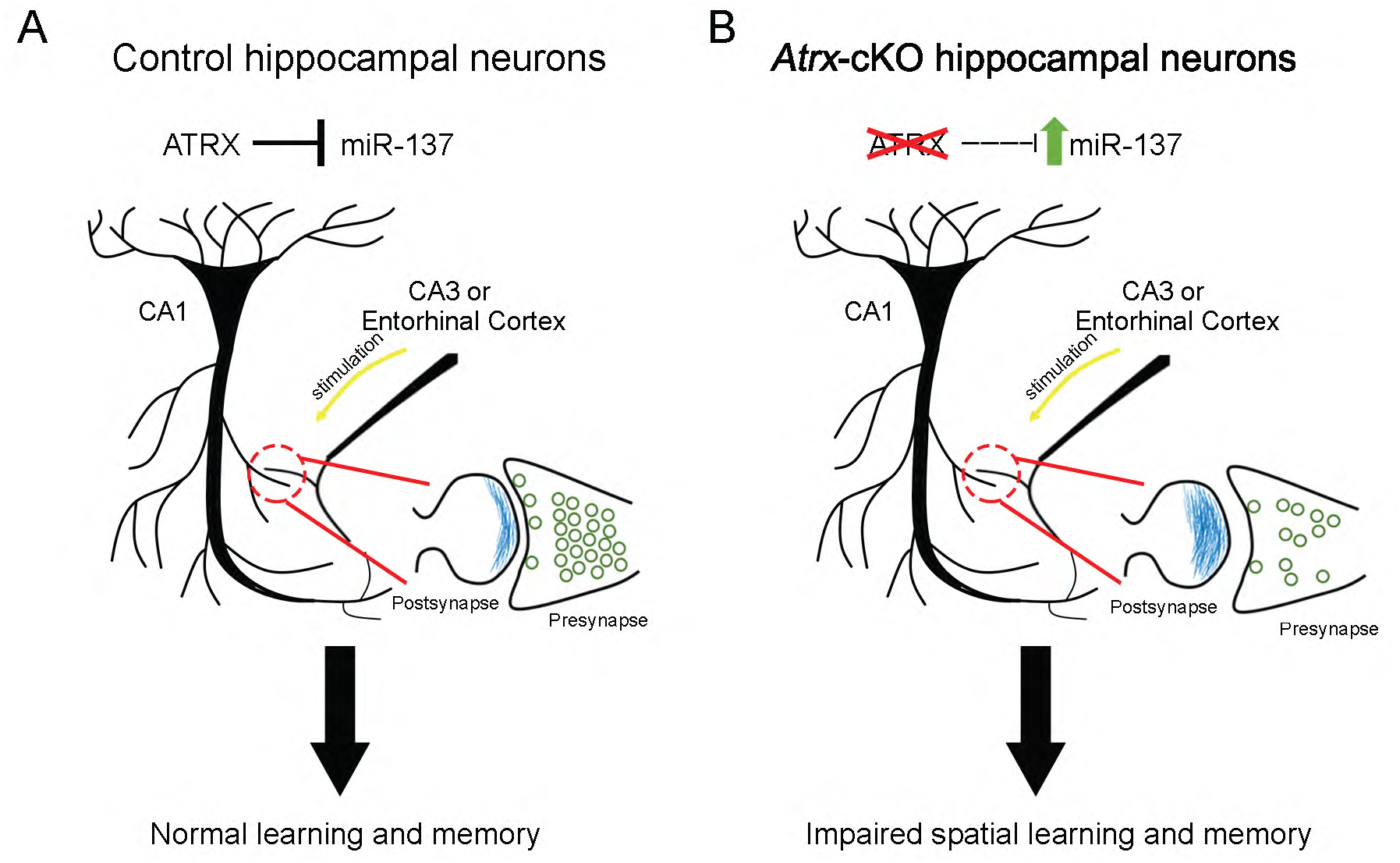
Proposed model of ATRX function in the hippocampus. (A) When present at normal levels, ATRX limits the expression of hippocampal miR-137 and alters target synaptic gene expression, thus preserving synaptic ultrastructural morphology and spatial learning and memory. (B) Loss of ATRX in forebrain glutamatergic neurons leads to increased hippocampal miR-137 expression in male mice, accompanied by ultrastructural synaptic defects, leading to sexually dimorphic defects in spatial learning and memory.

Loss of ATRX in the *Atrx*-cKO male mice resulted in various hippocampal-dependent memory impairments including contextual fear memory, and spatial learning and memory in the Morris water maze and paired-associate touchscreen task. The results from the touchscreen task were especially interesting considering the translational aspect of these tests^72^, and comparable results to humans are achieved using mouse models of cognitive impairments^51,73^. Unexpectedly, *Atrx*-cKO^Fem^ mice displayed normal memory in several paradigms, indicating a sex-specific phenomenon upon ATRX inactivation. In humans, females harbouring *ATRX* mutations are protected from the disease by complete skewing of X-inactivation^74^; however this cannot be the case here since the mice were homozygous for the “floxed” allele and we confirmed by immunofluorescence staining that ATRX is indeed absent from hippocampal excitatory neurons. Sexual dimorphism has been reported in other mouse models with mutations in chromatin remodeling proteins, including CHD8 and MeCP2, where females are unaffected by loss of the protein of interest or are affected differently^75-77^. In humans, neurological disorders such as autism-spectrum disorders tend to preferentially affect males rather than females, possibly due to combinatorial contributions of hormonal and genetic factors in a phenomenon known as the female protective effect^78-80^, and this is regularly supported with mouse models^81-83^. The presence of estrogen and estrogen receptor in the female brain has been shown to be neuroprotective and leads to enhanced Schaffer-collateral LTP^84^. In addition, certain X-linked genes involved in chromatin regulation (e.g. *Utx*, a histone demethylase) are able to escape X-inactivation and so are expressed two-fold in females compared to males^85^. These mechanisms could lead to protective gene regulation in the *Atrx*-cKO^Fem^, causing the sexually dimorphic defects in learning and memory. We previously reported impairment of spatial learning and memory in a female mouse model with mosaic expression of ATRX in all cells of the central nervous system^17^. This suggests the intriguing possibility that female-specific protective factors originate from cell types other than the excitatory neurons targeted in the present study. Female-specific glial factors, for example might provide important protection against the loss of ATRX in neurons.

MicroRNAs (miRNAs) are critical for the regulation of transcriptional programs in the brain. Excitation of whole neuronal networks results in drastic changes to miRNA levels, such as in seizures or traumatic brain injury^86,87^. miRNA levels can also change after exposure to certain behavioural tasks including contextual fear conditioning or novel object recognition^88,89^. At the molecular level, miRNAs can target 100s to 1000s of genes thereby altering expression of proteins at the presynaptic terminal, postsynaptic membrane, or both. The TEM and RNA-sequencing data obtained in the current study identified impaired presynaptic vesicular function, with decreased number of docked and total vesicles, prompting us to focus on miRNAs previously implicated in these functions: miR-485, miR-34a, miR-27b, and miR-137. Of the four miRNA tested, only miR-137 was significantly increased in *Atrx*-cKO male hippocampi while decreased in expression in the *Atrx*-cKO^Fem^ hippocampi, identifying another mechanism by which the females may be protected by ATRX loss. miR-137 overexpression has previously been linked to impaired hippocampal-dependent learning and memory in mice through the Morris water maze and contextual fear conditioning task, similar to the *Atrx*-cKO mice^58^. Additionally, miR-137 overexpression results in altered vesicular trafficking and reduced LTP *in vivo* and a reduction in the number of docked and total vesicles *in vitro*^*58*, *90*^, again paralleling our findings in the *Atrx*-cKO mice. miR-137 is also known to regulate expression of genes involved in postsynaptic function, including NMDA and AMPA receptor synthesis^91,92^ and multiple targets in the PI3K-Akt-mTOR pathway^93^. Therefore, increased expression of miR-137 could explain the presynaptic defects seen in the *Atrx-*cKO male mice and may have other effects within the post-synaptic density and downstream signalling that should be examined in more detail in the future.

The previously reported ATRX^ΔE2^ mice had impaired contextual fear memory, working memory, novel object recognition memory, and spatial memory in the Barnes maze^15,94^. Molecular analyses revealed decreased levels of CaMKII protein at synapses and increased expression of the *Xlr3b* gene, which is proposed to bind synaptic mRNAs and inhibit transport to dendritic spines^16^. In comparison, the *Atrx-*cKO male mice had no defects in working memory (Y-maze) or in novel object recognition. These differences point to potential cell-type specific effects, where ATRX downregulation or loss contributes to spatial memory, while loss in other cell types affects working and recognition memory. The timing of inactivation might also explain differences in behavioural outcomes. The ATRX^ΔE2^ mice may display a greater number of defects because ATRX depletion is present from the beginning of brain development, while *Atrx* inactivation occurs postnatally in the *CamKII*-Cre conditional approach taken here. Future investigations should discern the effects of inactivating ATRX in various cell types on cognitive output.

In conclusion, our study presents strong evidence that ATRX is required in forebrain excitatory neurons for spatial learning and long-term memory and regulation of genes required for efficient synaptic transmission.

## Acknowledgements

We are grateful to Doug Higgs and Richard Gibbons for the *Atrx* floxed mice, Vania Prado and Marco Prado for the *CaMKII-Cre* mice, Michael Miller for advice on statistical analyses, and Tim Bussey for discussions on the touchscreen assays. The dPAL and VPD touchscreen assays were performed at the Robarts Research Institute neurobehavioural core facility, TEM imaging at the Biotron at Western University and the RNA sequencing at the London Regional Genomics Centre.

## Author Contributions

R.J.T.: design, interpretation of data, execution of experiments, writing of article. V.D.: bioinformatics analyses. L.L.: execution of experiments. J.E. and L.R.Q.: execution of MRI imaging and data analysis. Y.J.: execution of experiments. J.L.: interpretation of data, editing the manuscript. N.G.B.: conception, design, interpretation of data, writing of article.

---

**Supplementary Figure 1:**
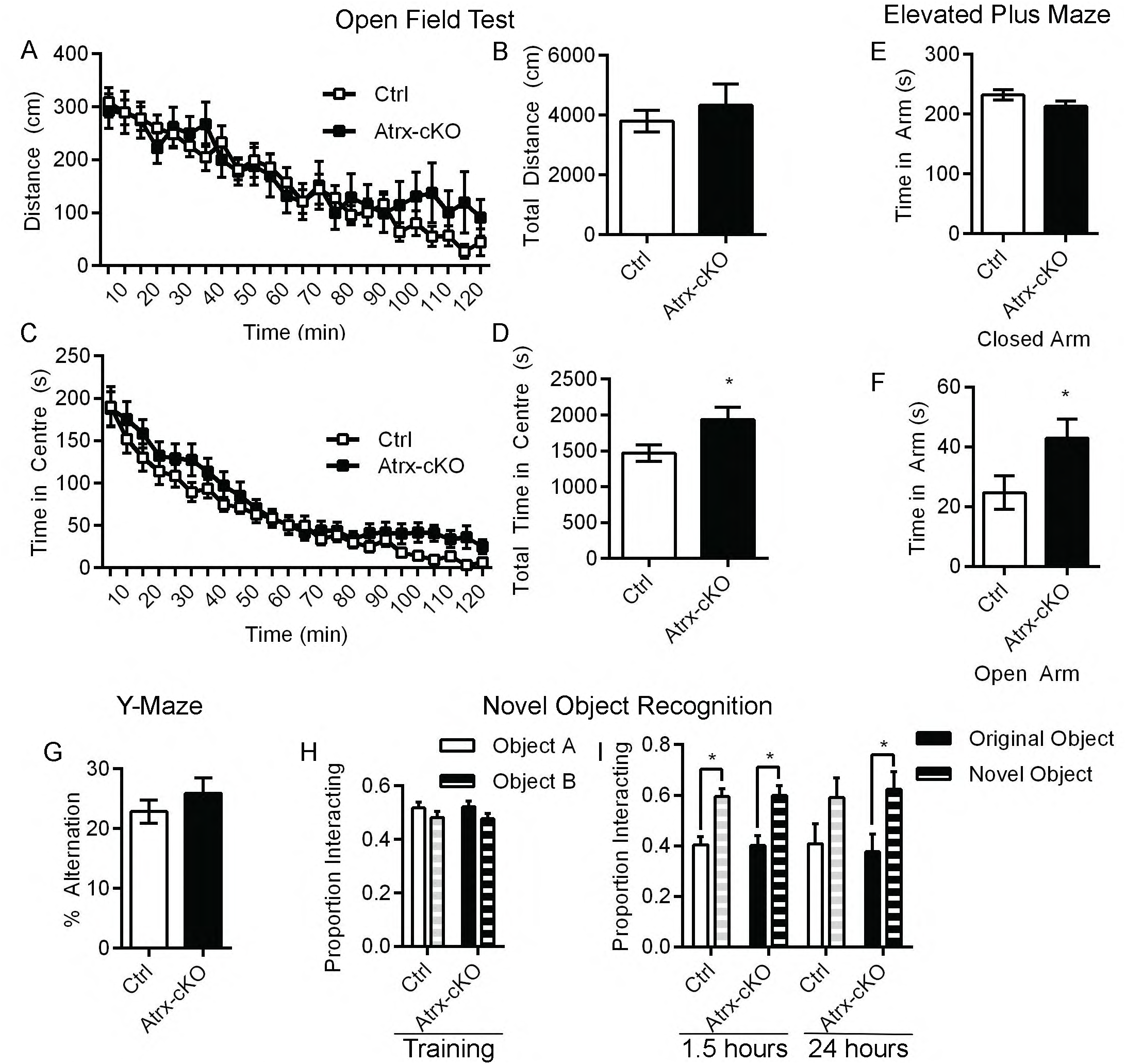
The *Atrx*-cKO males displayed decreased anxiety in the open field test and elevated plus maze. (A) Distance travelled over 120 minutes in the open field test (p=0.6803) (n=22). (B) Total distance travelled (p=0.5072). (C) Time spent in the centre over 120 minutes in the open field test (p=0.0927) (n=22). (D) Total time spent in the centre (p<0.05). (E) Time spent in the closed (p=0.1308) and open (p<0.05) arm of the elevated plus maze (n=14). Statistics by two-way repeated measures ANOVA with Sidak’s multiple comparisons test or Student’s T-test where applicable, asterisks indicate p<0.05.

**Supplementary Figure 2:**
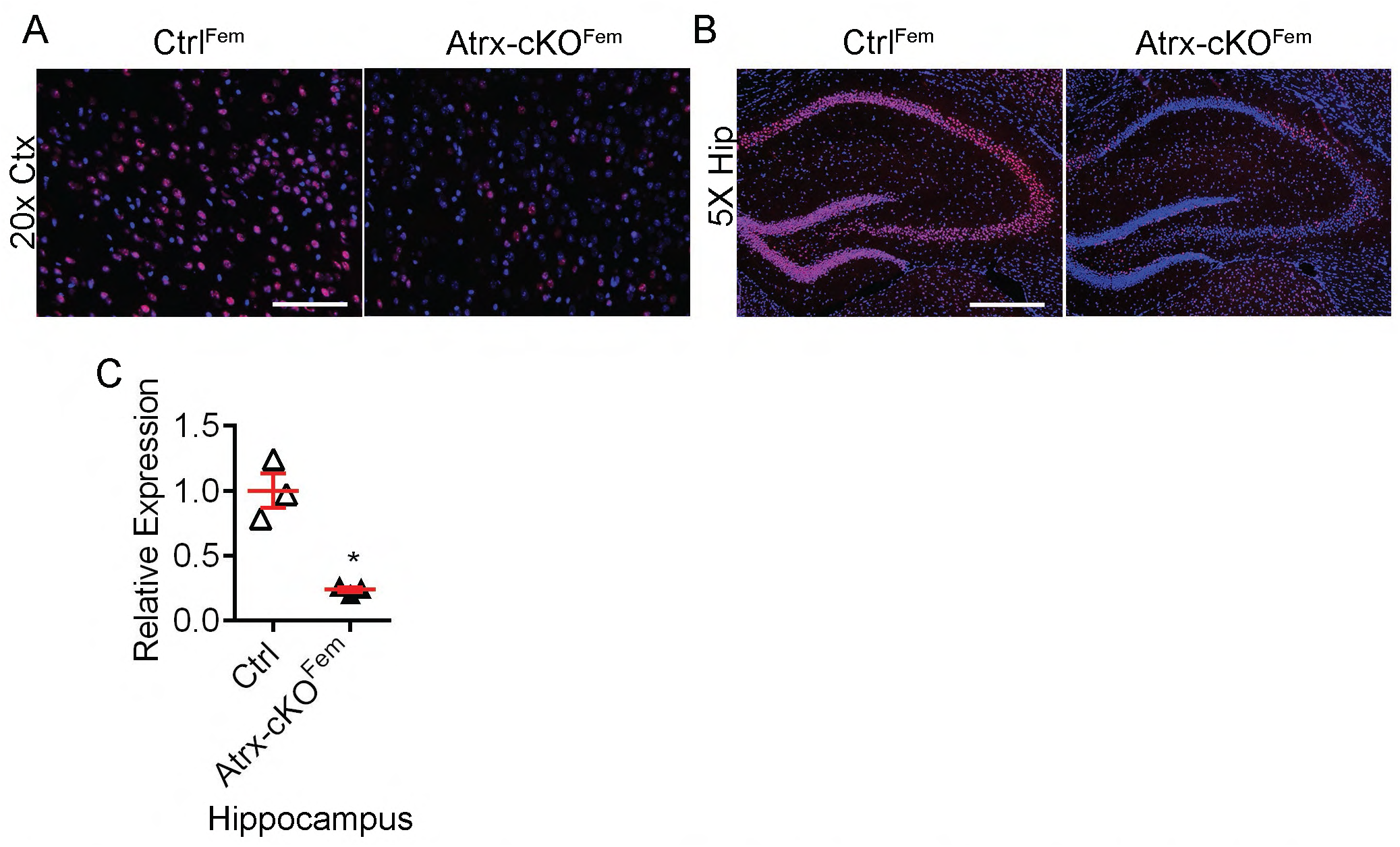
Expression of *Atrx* in *Atrx*-cKO females. (A) Immunofluorescence in cortex of control (Ctrl^Fem^) and knockout (*Atrx*-cKO^Fem^) female mice. ATRX: red; DAPI: blue. Scale bar: 50 µm. (B) Immunofluorescence in hippocampus of control (Ctrl^Fem^) and knockout (*Atrx*-cKO^Fem^) female mice. ATRX: red; DAPI: blue. Scale bar: 200 µm. (C) Expression of *Atrx* in the hippocampus by qRT-PCR (n=4) (p<0.005). Statistics by unpaired Student’s T-test.

**Supplementary Figure 3:**
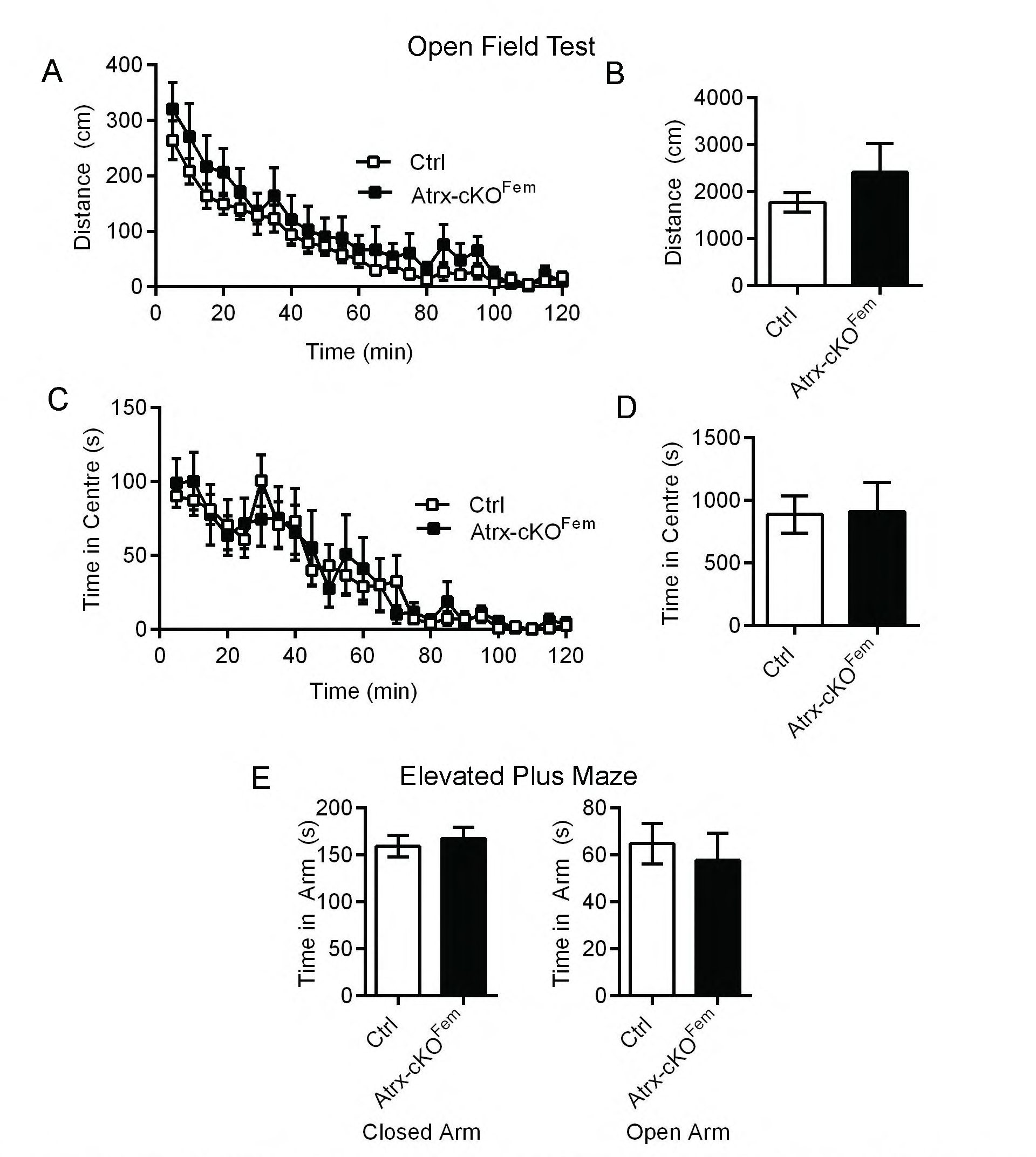
*Atrx*-cKO females have no change in anxiety. (A) Distance travelled over 120 minutes in the open field test (p=0.2796) (Ctrl^Fem^ n=12, *Atrx*-cKO^Fem^ n=9). (B) Total distance travelled (p=0.2796). (C) Time spent in the centre over 120 minutes in the open field test (p=0.9254) (Ctrl^Fem^ n=12, *Atrx*-cKO^Fem^ n=9). (D) Total time spent in the centre (p=0.9254). (E) Time spent in the closed (p=0.6277) and open (p=0.6250) arm of the elevated plus maze (Ctrl^Fem^ n=15, *Atrx*-cKO^Fem^ n=12). Statistics by two-way repeated measures ANOVA with Sidak’s multiple comparisons test or Student’s T-test where applicable.

**Supplementary Figure 4:**
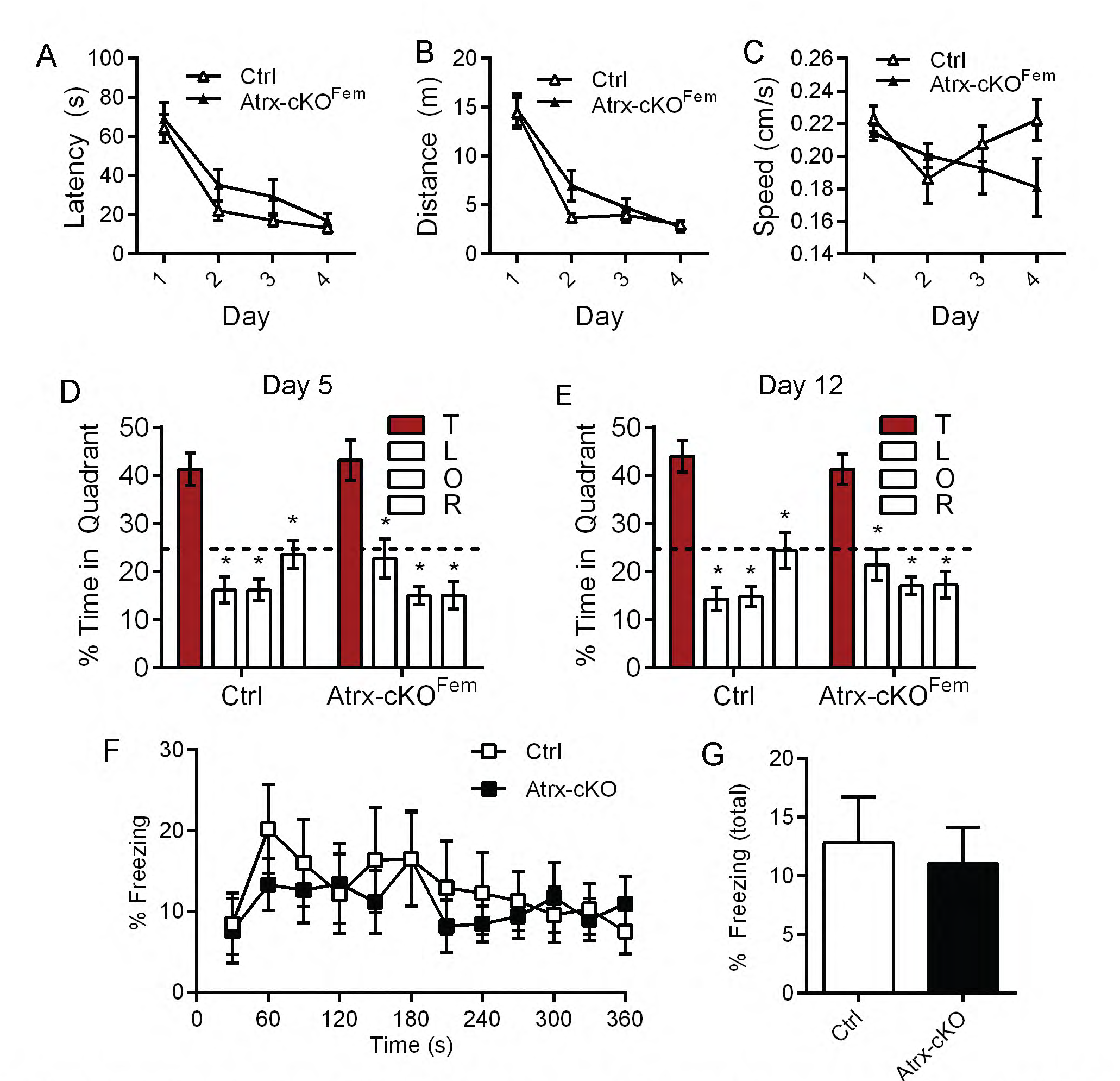
No change in spatial learning or memory in *Atrx*-cKO females in the Morris water maze or contextual fear conditioning tasks. (A,B,C) Latency (p=0.0683), distance (p=0.2503), and speed (p=0.2754) over four days (four trials per day) in the Morris water maze (Ctrl^Fem^ n=15, *Atrx*-cKO^Fem^ n=12). (D) Percent time spent in each quadrant after removal of the platform on day 5. Dotted line indicates chance at 25%. (E) Percent time spent in each quadrant after removal of the platform on day 12. Dotted line indicates chance at 25%. (F) Percent of time freezing over 360s during the contextual fear conditioning task (p=0.8818) (Ctrl^Fem^ n=15, *Atrx*-cKO^Fem^ n=12. (G) Total time freezing (p=0.8818). Statistics by two-way repeated measures ANOVA with Sidak’s multiple comparisons test or Student’s T-test where applicable, asterisks indicate p<0.05.

**Supplementary Figure 5:**
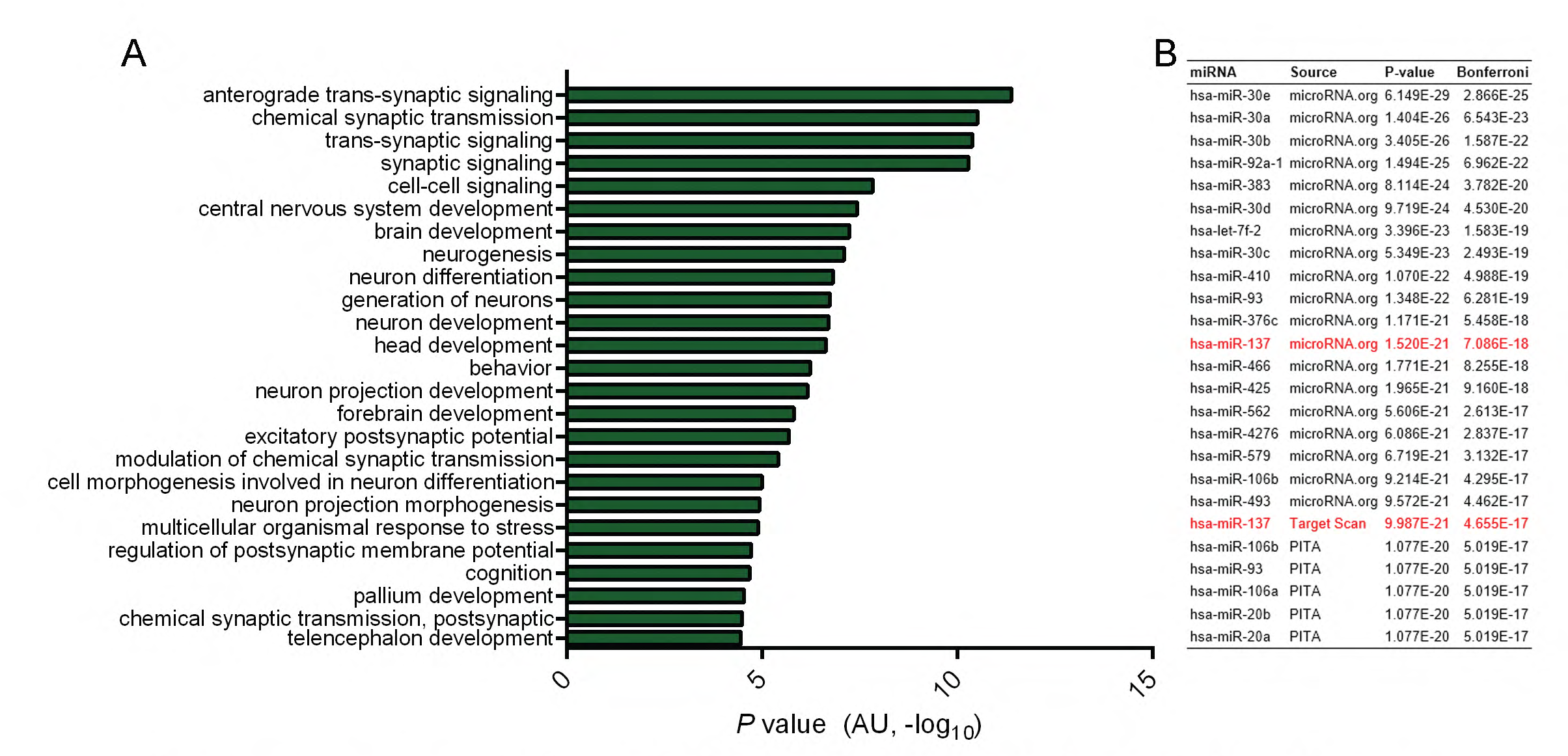
Transcriptional profiling reveals dysregulation of presynaptic genes in Atrx-FoxG1 mice. (A) Differentially expressed genes between Control and *Atrx*-FoxG1 P0.5 forebrain were used for Gene Ontology analysis and top 25 Biological Processes were listed by P-value. (B) Top miRNA predicted to regulate differentially expressed genes from Control and *Atrx*-FoxG1 mice.

## References

1. Gibbons, R. J., Picketts, D. J., Villard, L. & Higgs, D. R. Mutations in a putative global transcriptional regulator cause X-linked mental retardation with alpha-thalassemia (ATR-X syndrome). Cell 80, 837–845 (1995).

2. Grozeva, D. et al. Targeted Next-Generation Sequencing Analysis of 1,000 Individuals with Intellectual Disability. Human mutation 36, 1197–1204, doi:10.1002/humu.22901 (2015).

3. Gibbons, R. J. et al. Mutations in transcriptional regulator ATRX establish the functional significance of a PHD-like domain. Nature genetics 17, 146–148, doi:10.1038/ng1097-146 (1997).

4. Gibbons, R. J. et al. Mutations in the chromatin-associated protein ATRX. Human mutation 29, 796–802, doi:10.1002/humu.20734 (2008).

5. Eustermann, S. et al. Combinatorial readout of histone H3 modifications specifies localization of ATRX to heterochromatin. Nature structural & molecular biology 18, 777–782, doi:10.1038/nsmb.2070 (2011).

6. Picketts, D. J. et al. ATRX encodes a novel member of the SNF2 family of proteins: mutations point to a common mechanism underlying the ATR-X syndrome. Human molecular genetics 5, 1899–1907 (1996).

7. Lewis, P. W., Elsaesser, S. J., Noh, K. M., Stadler, S. C. & Allis, C. D. Daxx is an H3.3-specific histone chaperone and cooperates with ATRX in replication-independent chromatin assembly at telomeres. Proceedings of the National Academy of Sciences of the United States of America 107, 14075–14080, doi:10.1073/pnas.1008850107 (2010).

8. Goldberg, A. D. et al. Distinct factors control histone variant H3.3 localization at specific genomic regions. Cell 140, 678–691, doi:10.1016/j.cell.2010.01.003 (2010).

9. Law, M. J. et al. ATR-X syndrome protein targets tandem repeats and influences allele-specific expression in a size-dependent manner. Cell 143, 367–378, doi:10.1016/j.cell.2010.09.023 (2010).

10. Levy, M. A., Kernohan, K. D., Jiang, Y. & Berube, N. G. ATRX promotes gene expression by facilitating transcriptional elongation through guanine-rich coding regions. Human molecular genetics 24, 1824–1835, doi:10.1093/hmg/ddu596 (2015).

11. Kernohan, K. D. et al. ATRX partners with cohesin and MeCP2 and contributes to developmental silencing of imprinted genes in the brain. Developmental cell 18, 191–202, doi:10.1016/j.devcel.2009.12.017 (2010).

12. Garrick, D. et al. Loss of Atrx affects trophoblast development and the pattern of X-inactivation in extraembryonic tissues. PLoS genetics 2, e58, doi:10.1371/journal.pgen.0020058 (2006).

13. Watson, L. A. et al. Atrx deficiency induces telomere dysfunction, endocrine defects, and reduced life span. The Journal of clinical investigation 123, 2049–2063, doi:10.1172/JCI65634 (2013).

14. Seah, C. et al. Neuronal death resulting from targeted disruption of the Snf2 protein ATRX is mediated by p53. The Journal of neuroscience: the official journal of the Society for Neuroscience 28, 12570–12580, doi:10.1523/JNEUROSCI.4048-08.2008 (2008).

15. Shioda, N. et al. Aberrant calcium/calmodulin-dependent protein kinase II (CaMKII) activity is associated with abnormal dendritic spine morphology in the ATRX mutant mouse brain. The Journal of neuroscience: the official journal of the Society for Neuroscience 31, 346–358, doi:10.1523/JNEUROSCI.4816-10.2011 (2011).

16. Shioda, N. et al. Targeting G-quadruplex DNA as cognitive function therapy for ATR-X syndrome. Nature medicine 24, 802–813, doi:10.1038/s41591-018-0018-6 (2018).

17. Tamming, R. J. et al. Mosaic expression of Atrx in the mouse central nervous system causes memory deficits. Disease models & mechanisms 10, 119–126, doi:10.1242/dmm.027482 (2017).

18. Bérubé, N. G. et al. The chromatin-remodeling protein ATRX is critical for neuronal survival during corticogenesis. The Journal of clinical investigation 115, 258–267, doi:10.1172/JCI22329 (2005).

19. Tsien, J. Z. et al. Subregion- and cell type-restricted gene knockout in mouse brain. Cell 87, 1317-1326 (1996).

20. de Guzman, A. E., Wong, M. D., Gleave, J. A. & Nieman, B. J. Variations in post-perfusion immersion fixation and storage alter MRI measurements of mouse brain morphometry. NeuroImage 142, 687–695, doi:10.1016/j.neuroimage.2016.06.028 (2016).

21. Nieman, B. J. et al. MRI to Assess Neurological Function. Current protocols in mouse biology 8, e44, doi:10.1002/cpmo.44 (2018).

22. Spencer Noakes, T. L., Henkelman, R. M. & Nieman, B. J. Partitioning k-space for cylindrical three-dimensional rapid acquisition with relaxation enhancement imaging in the mouse brain. NMR in biomedicine 30, doi:10.1002/nbm.3802 (2017).

23. Collins, D. L., Neelin, P., Peters, T. M. & Evans, A. C. Automatic 3D intersubject registration of MR volumetric data in standardized Talairach space. Journal of computer assisted tomography 18, 192–205 (1994).

24. Avants, B. B., Tustison, N. J., Wu, J., Cook, P. A. & Gee, J. C. An open source multivariate framework for n-tissue segmentation with evaluation on public data. Neuroinformatics 9, 381–400, doi:10.1007/s12021-011-9109-y (2011).

25. Lerch, J. P. et al. Cortical thickness measured from MRI in the YAC128 mouse model of Huntington’s disease. NeuroImage 41, 243–251, doi:10.1016/j.neuroimage.2008.02.019 (2008).

26. Nieman, B. J., Flenniken, A. M., Adamson, S. L., Henkelman, R. M. & Sled, J. G. Anatomical phenotyping in the brain and skull of a mutant mouse by magnetic resonance imaging and computed tomography. Physiological genomics 24, 154–162, doi:10.1152/physiolgenomics.00217.2005 (2006).

27. Qiu, L. R. et al. Mouse MRI shows brain areas relatively larger in males emerge before those larger in females. Nature communications 9, 2615, doi:10.1038/s41467-018-04921-2 (2018).

28. Dorr, A. E., Lerch, J. P., Spring, S., Kabani, N. & Henkelman, R. M. High resolution three-dimensional brain atlas using an average magnetic resonance image of 40 adult C57Bl/6J mice. NeuroImage 42, 60–69, doi:10.1016/j.neuroimage.2008.03.037 (2008).

29. Steadman, P. E. et al. Genetic effects on cerebellar structure across mouse models of autism using a magnetic resonance imaging atlas. Autism research: official journal of the International Society for Autism Research 7, 124–137, doi:10.1002/aur.1344 (2014).

30. Ullmann, J. F. et al. An MRI atlas of the mouse basal ganglia. Brain structure & function 219, 1343–1353, doi:10.1007/s00429-013-0572-0 (2014).

31. Richards, K. et al. Segmentation of the mouse hippocampal formation in magnetic resonance images. NeuroImage 58, 732–740, doi:10.1016/j.neuroimage.2011.06.025 (2011).

32. Genovese, C. R., Lazar, N. A. & Nichols, T. Thresholding of statistical maps in functional neuroimaging using the false discovery rate. NeuroImage 15, 870–878, doi:10.1006/nimg.2001.1037 (2002).

33. Longair, M. H., Baker, D. A. & Armstrong, J. D. Simple Neurite Tracer: open source software for reconstruction, visualization and analysis of neuronal processes. Bioinformatics 27, 2453–2454, doi:10.1093/bioinformatics/btr390 (2011).

34. Ferreira, T. A. et al. Neuronal morphometry directly from bitmap images. Nature methods 11, 982–984, doi:10.1038/nmeth.3125 (2014).

35. de Castro, B. M. et al. Reduced expression of the vesicular acetylcholine transporter causes learning deficits in mice. Genes, brain, and behavior 8, 23–35, doi:10.1111/j.1601-183X.2008.00439.x (2009).

36. Vorhees, C. V. & Williams, M. T. Morris water maze: procedures for assessing spatial and related forms of learning and memory. Nature protocols 1, 848–858, doi:10.1038/nprot.2006.116 (2006).

37. Talpos, J. C., Winters, B. D., Dias, R., Saksida, L. M. & Bussey, T. J. A novel touchscreen-automated paired-associate learning (PAL) task sensitive to pharmacological manipulation of the hippocampus: a translational rodent model of cognitive impairments in neurodegenerative disease. Psychopharmacology 205, 157–168, doi:10.1007/s00213-009-1526-3 (2009).

38. Bussey, T. J. et al. The touchscreen cognitive testing method for rodents: how to get the best out of your rat. Learning & memory 15, 516–523, doi:10.1101/lm.987808 (2008).

39. Delotterie, D. F. et al. Touchscreen tasks in mice to demonstrate differences between hippocampal and striatal functions. Neurobiology of learning and memory 120, 16–27, doi:10.1016/j.nlm.2015.02.007 (2015).

40. Zerbino, D. R. et al. Ensembl 2018. Nucleic acids research 46, D754-D761, doi:10.1093/nar/gkx1098 (2018).

41. Pertea, M. et al. StringTie enables improved reconstruction of a transcriptome from RNA-seq reads. Nature biotechnology 33, 290–295, doi:10.1038/nbt.3122 (2015).

42. Soneson, C., Love, M. I. & Robinson, M. D. Differential analyses for RNA-seq: transcript-level estimates improve gene-level inferences. F1000Research 4, 1521, doi:10.12688/f1000research.7563.2 (2015).

43. Love, M. I., Huber, W. & Anders, S. Moderated estimation of fold change and dispersion for RNA-seq data with DESeq2. Genome biology 15, 550, doi:10.1186/s13059-014-0550-8 (2014).

44. Ignatiadis, N., Klaus, B., Zaugg, J. B. & Huber, W. Data-driven hypothesis weighting increases detection power in genome-scale multiple testing. Nature methods 13, 577–580, doi:10.1038/nmeth.3885 (2016).

45. Benjamini, Y. & Hochberg, Y. Controlling the false discovery rate: a practical and powerful approach to multiple testing. Journal of the Royal statistical society: series B (Methodological) 57, 289–300 (1995).

46. Alexa, A., Rahnenfuhrer, J. & Lengauer, T. Improved scoring of functional groups from gene expression data by decorrelating GO graph structure. Bioinformatics 22, 1600–1607, doi:10.1093/bioinformatics/btl140 (2006).

47. Harris, K. M., Jensen, F. E. & Tsao, B. Three-dimensional structure of dendritic spines and synapses in rat hippocampus (CA1) at postnatal day 15 and adult ages: implications for the maturation of synaptic physiology and long-term potentiation. The Journal of neuroscience: the official journal of the Society for Neuroscience 12, 2685–2705 (1992).

48. Bisht, K., El Hajj, H., Savage, J. C., Sanchez, M. G. & Tremblay, M. E. Correlative Light and Electron Microscopy to Study Microglial Interactions with beta-Amyloid Plaques. Journal of visualized experiments: JoVE, doi:10.3791/54060 (2016).

49. Tronche, F. et al. Disruption of the glucocorticoid receptor gene in the nervous system results in reduced anxiety. Nature genetics 23, 99–103, doi:10.1038/12703 (1999).

50. Sahakian, B. J. et al. A comparative study of visuospatial memory and learning in Alzheimer-type dementia and Parkinson’s disease. Brain: a journal of neurology 111 (Pt 3), 695–718 (1988).

51. Nithianantharajah, J. et al. Bridging the translational divide: identical cognitive touchscreen testing in mice and humans carrying mutations in a disease-relevant homologous gene. Scientific reports 5, 14613, doi:10.1038/srep14613 (2015).

52. Kim, C. H., Heath, C. J., Kent, B. A., Bussey, T. J. & Saksida, L. M. The role of the dorsal hippocampus in two versions of the touchscreen automated paired associates learning (PAL) task for mice. Psychopharmacology 232, 3899–3910, doi:10.1007/s00213-015-3949-3 (2015).

53. Mi, H. et al. PANTHER version 11: expanded annotation data from Gene Ontology and Reactome pathways, and data analysis tool enhancements. Nucleic acids research 45, D183–D189, doi:10.1093/nar/gkw1138 (2017).

54. Ryan, B., Joilin, G. & Williams, J. M. Plasticity-related microRNA and their potential contribution to the maintenance of long-term potentiation. Frontiers in molecular neuroscience 8, 4, doi:10.3389/fnmol.2015.00004 (2015).

55. Cohen, J. E., Lee, P. R., Chen, S., Li, W. & Fields, R. D. MicroRNA regulation of homeostatic synaptic plasticity. Proceedings of the National Academy of Sciences of the United States of America 108, 11650–11655, doi:10.1073/pnas.1017576108 (2011).

56. Agostini, M. et al. microRNA-34a regulates neurite outgrowth, spinal morphology, and function. Proceedings of the National Academy of Sciences of the United States of America 108, 21099–21104, doi:10.1073/pnas.1112063108 (2011).

57. Poon, V. Y., Gu, M., Ji, F., VanDongen, A. M. & Fivaz, M. miR-27b shapes the presynaptic transcriptome and influences neurotransmission by silencing the polycomb group protein Bmi1. BMC genomics 17, 777, doi:10.1186/s12864-016-3139-7 (2016).

58. Siegert, S. et al. The schizophrenia risk gene product miR-137 alters presynaptic plasticity. Nature neuroscience 18, 1008–1016, doi:10.1038/nn.4023 (2015).

59. Levy, M. A., Fernandes, A. D., Tremblay, D. C., Seah, C. & Berube, N. G. The SWI/SNF protein ATRX co-regulates pseudoautosomal genes that have translocated to autosomes in the mouse genome. BMC genomics 9, 468, doi:10.1186/1471-2164-9-468 (2008).

60. Chen, J., Bardes, E. E., Aronow, B. J. & Jegga, A. G. ToppGene Suite for gene list enrichment analysis and candidate gene prioritization. Nucleic acids research 37, W305-311, doi:10.1093/nar/gkp427 (2009).

61. Sheng, M. & Kim, E. The Shank family of scaffold proteins. Journal of cell science 113 (Pt 11), 1851–1856 (2000).

62. Rajendra, S., Lynch, J. W. & Schofield, P. R. The glycine receptor. Pharmacology & therapeutics 73, 121–146 (1997).

63. Cisternas, F. A., Vincent, J. B., Scherer, S. W. & Ray, P. N. Cloning and characterization of human CADPS and CADPS2, new members of the Ca2+-dependent activator for secretion protein family. Genomics 81, 279–291 (2003).

64. Dergai, O. et al. Intersectin 1 forms complexes with SGIP1 and Reps1 in clathrin-coated pits. Biochemical and biophysical research communications 402, 408–413, doi:10.1016/j.bbrc.2010.10.045 (2010).

65. Vago, D. R. & Kesner, R. P. Disruption of the direct perforant path input to the CA1 subregion of the dorsal hippocampus interferes with spatial working memory and novelty detection. Behavioural brain research 189, 273–283, doi:10.1016/j.bbr.2008.01.002 (2008).

66. Nguyen, P. V. & Kandel, E. R. A macromolecular synthesis-dependent late phase of long-term potentiation requiring cAMP in the medial perforant pathway of rat hippocampal slices. The Journal of neuroscience: the official journal of the Society for Neuroscience 16, 3189–3198 (1996).

67. McGill, B. E. et al. Abnormal Microglia and Enhanced Inflammation-Related Gene Transcription in Mice with Conditional Deletion of Ctcf in Camk2a-Cre-Expressing Neurons. The Journal of neuroscience: the official journal of the Society for Neuroscience 38, 200–219, doi:10.1523/JNEUROSCI.0936-17.2017 (2018).

68. Tanaka, S. et al. Lipopolysaccharide-induced microglial activation induces learning and memory deficits without neuronal cell death in rats. Journal of neuroscience research 83, 557–566, doi:10.1002/jnr.20752 (2006).

69. Bian, Y. et al. Learning, memory, and glial cell changes following recovery from chronic unpredictable stress. Brain research bulletin 88, 471–476, doi:10.1016/j.brainresbull.2012.04.008 (2012).

70. Hylin, M. J., Orsi, S. A., Moore, A. N. & Dash, P. K. Disruption of the perineuronal net in the hippocampus or medial prefrontal cortex impairs fear conditioning. Learning & memory 20, 267–273, doi:10.1101/lm.030197.112 (2013).

71. Bukalo, O., Schachner, M. & Dityatev, A. Modification of extracellular matrix by enzymatic removal of chondroitin sulfate and by lack of tenascin-R differentially affects several forms of synaptic plasticity in the hippocampus. Neuroscience 104, 359–369 (2001).

72. Bussey, T. J. et al. New translational assays for preclinical modelling of cognition in schizophrenia: the touchscreen testing method for mice and rats. Neuropharmacology 62, 1191–1203, doi:10.1016/j.neuropharm.2011.04.011 (2012).

73. Nithianantharajah, J. et al. Synaptic scaffold evolution generated components of vertebrate cognitive complexity. Nature neuroscience 16, 16–24, doi:10.1038/nn.3276 (2013).

74. Gibbons, R. J., Suthers, G. K., Wilkie, A. O., Buckle, V. J. & Higgs, D. R. X-linked alpha-thalassemia/mental retardation (ATR-X) syndrome: localization to Xq12-q21.31 by X inactivation and linkage analysis. American journal of human genetics 51, 1136–1149 (1992).

75. Jung, H. et al. Sexually dimorphic behavior, neuronal activity, and gene expression in Chd8-mutant mice. Nature neuroscience 21, 1218–1228, doi:10.1038/s41593-018-0208-z (2018).

76. Kurian, J. R., Bychowski, M. E., Forbes-Lorman, R. M., Auger, C. J. & Auger, A. P. Mecp2 organizes juvenile social behavior in a sex-specific manner. The Journal of neuroscience: the official journal of the Society for Neuroscience 28, 7137–7142, doi:10.1523/JNEUROSCI.1345-08.2008 (2008).

77. Kim, K. C. et al. MeCP2 Modulates Sex Differences in the Postsynaptic Development of the Valproate Animal Model of Autism. Molecular neurobiology 53, 40–56, doi:10.1007/s12035-014-8987-z (2016).

78. Fombonne, E. Epidemiology of pervasive developmental disorders. Pediatric research 65, 591–598, doi:10.1203/PDR.0b013e31819e7203 (2009).

79. Jacquemont, S. et al. A higher mutational burden in females supports a “female protective model” in neurodevelopmental disorders. American journal of human genetics 94, 415–425, doi:10.1016/j.ajhg.2014.02.001 (2014).

80. Voineagu, I. et al. Transcriptomic analysis of autistic brain reveals convergent molecular pathology. Nature 474, 380–384, doi:10.1038/nature10110 (2011).

81. Tsutiya, A. et al. Human CRMP4 mutation and disrupted Crmp4 expression in mice are associated with ASD characteristics and sexual dimorphism. Scientific reports 7, 16812, doi:10.1038/s41598-017-16782-8 (2017).

82. Sato, D. et al. SHANK1 Deletions in Males with Autism Spectrum Disorder. American journal of human genetics 90, 879–887, doi:10.1016/j.ajhg.2012.03.017 (2012).

83. Hu, V. W., Sarachana, T., Sherrard, R. M. & Kocher, K. M. Investigation of sex differences in the expression of RORA and its transcriptional targets in the brain as a potential contributor to the sex bias in autism. Molecular autism 6, 7, doi:10.1186/2040-2392-6-7 (2015).

84. Wang, W. et al. Memory-Related Synaptic Plasticity Is Sexually Dimorphic in Rodent Hippocampus. The Journal of neuroscience: the official journal of the Society for Neuroscience 38, 7935–7951, doi:10.1523/JNEUROSCI.0801-18.2018 (2018).

85. Xu, J., Deng, X., Watkins, R. & Disteche, C. M. Sex-specific differences in expression of histone demethylases Utx and Uty in mouse brain and neurons. The Journal of neuroscience: the official journal of the Society for Neuroscience 28, 4521–4527, doi:10.1523/JNEUROSCI.5382-07.2008 (2008).

86. Jimenez-Mateos, E. M. et al. miRNA Expression profile after status epilepticus and hippocampal neuroprotection by targeting miR-132. The American journal of pathology 179, 2519–2532, doi:10.1016/j.ajpath.2011.07.036 (2011).

87. Lei, P., Li, Y., Chen, X., Yang, S. & Zhang, J. Microarray based analysis of microRNA expression in rat cerebral cortex after traumatic brain injury. Brain research 1284, 191–201, doi:10.1016/j.brainres.2009.05.074 (2009).

88. Lin, Q. et al. The brain-specific microRNA miR-128b regulates the formation of fear-extinction memory. Nature neuroscience 14, 1115–1117, doi:10.1038/nn.2891 (2011).

89. Woldemichael, B. T. et al. The microRNA cluster miR-183/96/182 contributes to long-term memory in a protein phosphatase 1-dependent manner. Nature communications 7, 12594, doi:10.1038/ncomms12594 (2016).

90. He, E. et al. MIR137 schizophrenia-associated locus controls synaptic function by regulating synaptogenesis, synapse maturation and synaptic transmission. Human molecular genetics 27, 1879–1891, doi:10.1093/hmg/ddy089 (2018).

91. Zhao, L. et al. miR-137, a new target for post-stroke depression? Neural regeneration research 8, 2441–2448, doi:10.3969/j.issn.1673-5374.2013.26.005 (2013).

92. Olde Loohuis, N. F. et al. MicroRNA-137 Controls AMPA-Receptor-Mediated Transmission and mGluR-Dependent LTD. Cell reports 11, 1876–1884, doi:10.1016/j.celrep.2015.05.040 (2015).

93. Thomas, K. T. et al. Inhibition of the Schizophrenia-Associated MicroRNA miR-137 Disrupts Nrg1alpha Neurodevelopmental Signal Transduction. Cell reports 20, 1–12, doi:10.1016/j.celrep.2017.06.038 (2017).

94. Nogami, T. et al. Reduced expression of the ATRX gene, a chromatin-remodeling factor, causes hippocampal dysfunction in mice. Hippocampus 21, 678–687, doi:10.1002/hipo.20782 (2011).

